# Visualizing a two-state conformational ensemble in stem-loop 3 of the transcriptional regulator 7SK RNA

**DOI:** 10.1101/2023.08.09.552709

**Authors:** Momodou B. Camara, Bret Lange, Joseph D. Yesselman, Catherine D. Eichhorn

**Affiliations:** Department of Chemistry, University of Nebraska, 639 North 12th St, Lincoln, NE 68588, USA; Nebraska Center for Integrated Biomolecular Communication

**Keywords:** solution NMR spectroscopy, conformational exchange, chemical mapping, RNA dynamics, phylogenetic analysis

## Abstract

Structural plasticity is integral to RNA function; however, there are currently few methods to quantitatively resolve RNAs that have multiple structural states. NMR spectroscopy is a powerful approach for resolving conformational ensembles but is size-limited. Chemical probing is well-suited for large RNAs but provides limited structural and no kinetics information. Here, we integrate the two approaches to visualize a two-state conformational ensemble for the central stem-loop 3 (SL3) of 7SK RNA, a critical element for 7SK RNA function in transcription regulation. We find that the SL3 distal end exchanges between two equally populated yet structurally distinct states in both isolated SL3 constructs and full-length 7SK RNA. We rationally designed constructs that lock SL3 into a single state and demonstrate that both chemical probing and NMR data fit to a linear combination of the two states. Comparison of vertebrate 7SK RNA sequences shows conservation of both states, suggesting functional importance. These results provide new insights into 7SK RNA structural dynamics and demonstrate the utility of integrating chemical probing with NMR spectroscopy to gain quantitative insights into RNA conformational ensembles.

## Introduction

RNA secondary structure rearrangements are critical for cellular function^1–4^; however, there are at present few methods to quantitatively resolve these dynamic ensembles. NMR spectroscopy is a powerful method for resolving conformational ensembles at atomic resolution^2,5–7^ but can be challenging for large RNAs. In contrast chemical probing, particularly mutational profiling (MaP)-based chemical probing and next generation sequencing, is a powerful technique for identifying and deconvoluting conformational ensembles in large RNAs^8–12^. Many MaP techniques have been developed^4^; however, few of the identified RNA conformational ensembles have been validated by orthogonal approaches. At present, MaP techniques cannot measure kinetics and are limited in their ability to resolve local differences in structure^4,10^.

The noncoding 7SK RNA assembles with proteins to form the 7SK ribonucleoprotein (RNP) to negatively regulate eukaryotic transcription elongation through sequestration and inactivation of the positive transcription elongation factor b (P-TEFb)^13–17^. The emerging mechanism of P-TEFb release involves large-scale structural remodeling of 7SK RNA, which is proposed to be driven through protein association^11,18,19^. 7SK RNA folds into four primary stem-loop (SL) domains (**Fig 1a**). Several 7SK RNA secondary structure models have been reported from chemical and enzymatic probing or bioinformatics techniques^11,13,15,18–23^. In particular, multiple secondary structures have been proposed for the distal end of SL3, an element required to recruit accessory proteins to promote P-TEFb release on specific cellular cues^13,19,24–28^. Given the functional significance of SL3, there is an urgent need to characterize SL3 structural features to resolve these discrepancies.

**Figure 1.**
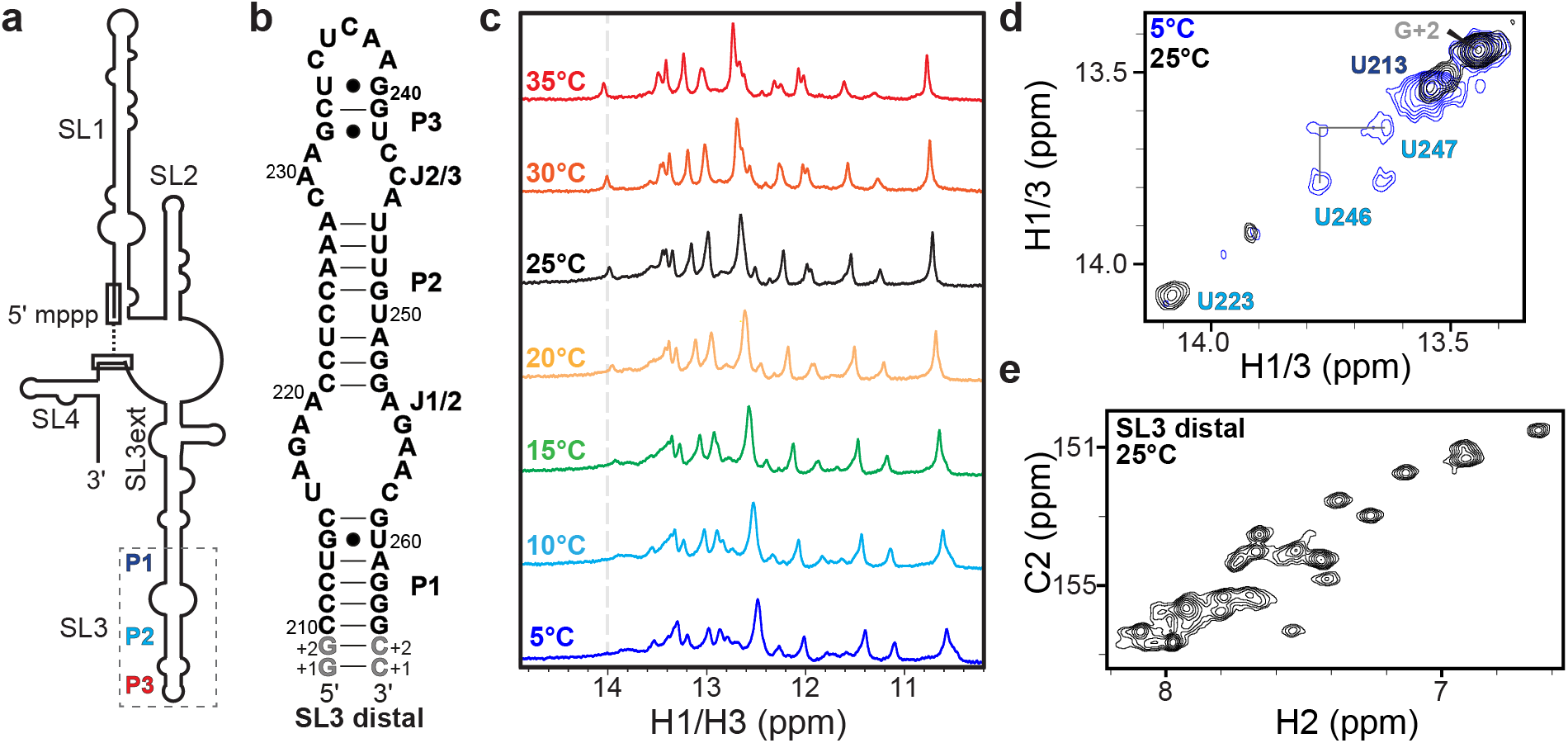
7SK RNA SL3 construct design and NMR evidence of conformational exchange a) Cartoon schematic of 7SK RNA secondary structure; b) SL3 distal domain (nts 210-264) constructs for NMR studies; c) 1D ^1^H imino proton NMR spectra show new resonances appear at elevated temperatures; d) 2D ^1^H-^1^H NOESY spectra show differences at 5 °C and 25 °C; e) 2D ^1^H-^13^C HSQC spectrum of adenine C2H2 resonances shows greater than the expected number of resonances indicative of slow exchange between multiple states.

Here we use solution NMR spectroscopy, optical melting, and dimethyl sulfate mutational profiling with sequencing (DMS-MaPseq) chemical probing approaches to characterize the distal end of SL3. We find that SL3 slowly exchanges between two nearly equally populated secondary structural states. This two-state exchange is present in SL3 of an isolated domain construct as well as *in vitro* full-length 7SK RNA. Using rational design, we generate subdomain and point mutation constructs that lock SL3 into individual states. We demonstrate that the SL3 distal DMS-MaPseq data can be explained by a linear combination of the two states, providing strong evidence that these secondary structures are in fact populated. A survey of vertebrate 7SK RNA SL3 sequences shows broad conservation of both states, provides insights into the sequence determinants for two-state exchange equilibrium, and suggests a functional role. Our results reconcile disparities in the literature by showing that rather than a single state, SL3 exists as an ensemble of two secondary structural states. Further, these studies demonstrate the utility in applying combined NMR and chemical probing approaches to quantitatively resolve RNA conformational ensembles.

## Results

### NMR reveals that the 7SK RNA SL3 distal domain adopts multiple conformations

We used solution NMR spectroscopy to characterize a domain construct of the SL3 distal sequence of human 7SK RNA (SL3 distal, nts 210-264) (**Fig. 1b**). The 1D ^1^H spectrum showed more imino proton resonances than expected, as well as temperature-dependent line broadening, indicating base-pair related chemical exchange (**Fig. 1c**). Due to the observed chemical exchange, the secondary structure could not be unambiguously determined from ^1^H-^1^H NOESY experiments. Imino proton resonances for P1 and P3 stem residues could be assigned from ^1^H-^1^H NOESY experiments, although P3 stem imino proton resonances were only observed at reduced temperatures (**Supplementary** Fig. 1). In contrast, few NOEs were observed for imino proton resonances attributed to the P2 stem (**Fig. 1d, Supplementary Fig. S1**). Consistent with 1D ^1^H NMR spectra, the ^1^H-^1^H NOESY spectra of the imino proton region showed a prominent on-diagonal peak at 14.1 ppm (previously assigned as U223^20^) at 25 °C but not 5 °C (**Fig. 1d**). Located in the middle of the P2 stem, U223 is not expected to undergo chemical exchange. We prepared a ^13^C/^15^N isotopically labeled sample and observed a greater-than-expected number of resonances in ^1^H-^13^C and ^1^H-^15^N HSQC spectra (**Supplementary Fig. S1**). For example, although the SL3 distal construct has 16 adenine residues over 20 C2H2 (adenine) resonances were observed (**Fig. 1e**). A possible explanation for this observation is that the RNA adopts more than one conformation that interconvert slowly on the NMR chemical shift timescale (typically millisecond or slower^29^). The imino proton resonance line broadening for P2 stem residues, and extra nucleobase resonances, indicates that SL3 has conformational exchange on slow exchange timescales.

### DMS-MaPseq shows 7SK SL3 adopts two conformations with distinct secondary structures

As an orthogonal approach to investigate the secondary structure, we performed DMS-MaPseq^10^ experiments on the isolated SL3 distal construct. Due to the lack of commercial availability of TGIRT-III enzyme, both TGIRT-III and Marathon reverse transcriptases were used with excellent agreement, as seen in prior studies^30,31^ (**Supplementary Fig. S2**). The secondary structure, assuming a single state, matched the secondary structure shown in **Fig. 1b**. However, P2 stem residues C221 and A251 had intermediate mutational fraction values, suggesting that a single secondary structure may be insufficient to describe the data (**Fig. 2a**). Next, we applied DREEM (Detection of RNA folding Ensembles using Expectation-Maximization)^32^ to investigate whether the DMS-MaPseq data could be deconvoluted into two or more secondary structure states. From DREEM, two clusters were identified with near-equal populations (**Fig. 2b-d**). Both clusters have identical P1 and P3 stems but differ significantly in the P2 stem and adjacent loops. A comparison of the mutational fraction values for the two clusters identified several key reporter residues in the center of the P2 stem and J1/2 and J2/3 loops that are present in a stem in one state and loop in the other state (**Fig. 2d**).

**Figure 2.**
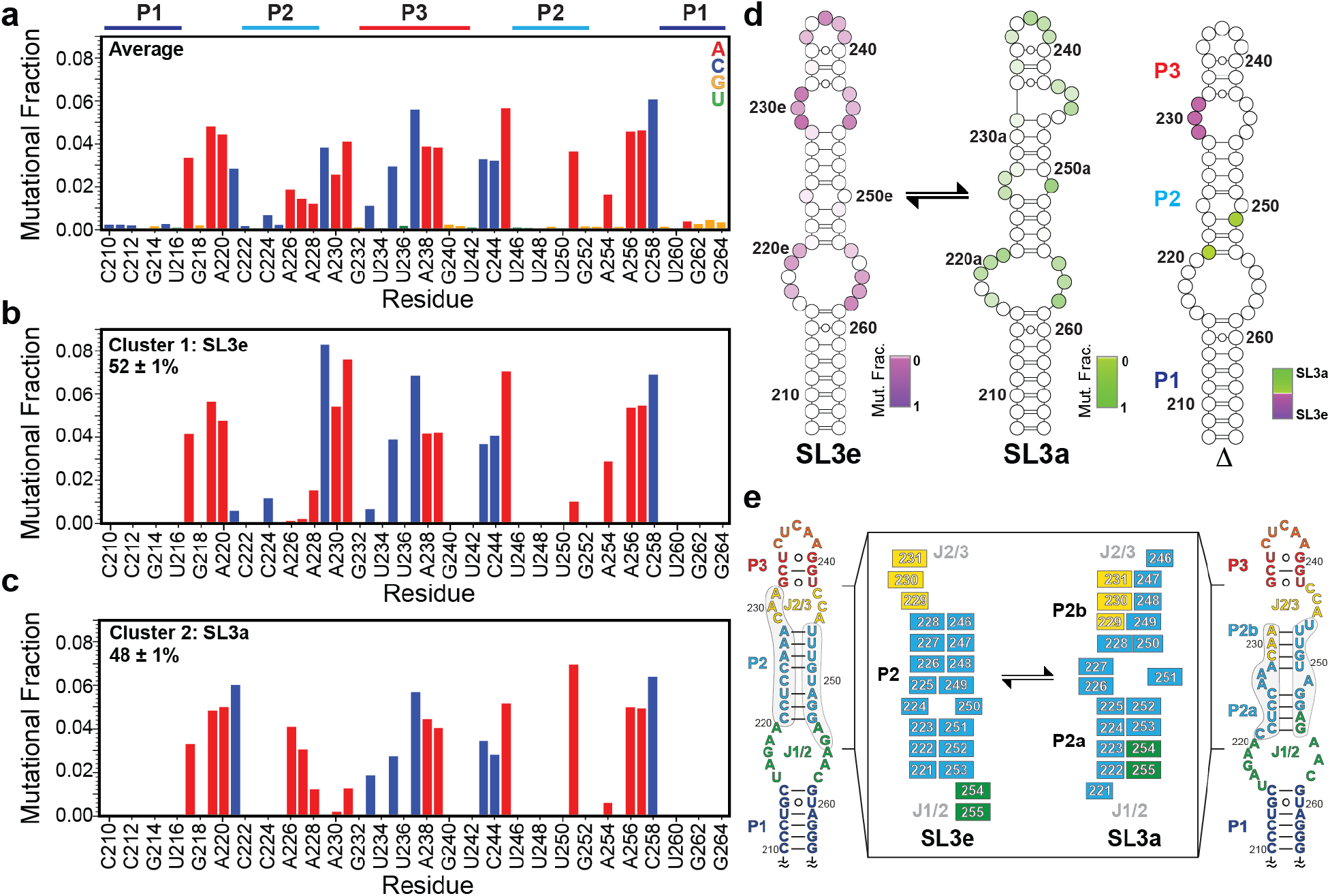
DMS-MaPseq and DREEM clustering supports the presence of two conformers. a) Average mutational fraction of SL3 distal construct. Residues are colored by nucleotide (A, red; C, blue; G, yellow; U, green). b-c) mutational fraction after DREEM clustering. d) Secondary structure models of SL3e (left), SL3a (center), and difference plot (right) colored according to DMS-MaPseq reactivity shows key reporter residues in the P2 stem and J2/J3 loop. e) Secondary structure model of SL3e, colored by motif (P1 dark blue, J1/2 green, P2 light blue, J2/3 yellow, P3 red). Inset: Schematic of differences between SL3e and SL3a secondary structures

The P2 stem of one cluster matched the secondary structure in **Fig. 1b**, hereafter named SL3e for an extended P2 stem (**Fig. 2d-e**). The P2 stem of SL3e has eight base-pairs with a central C•U mismatch. This secondary structure was observed in prior chemical probing and NMR studies^11,20^. The P2 stem of the other cluster consists of two four base-pair stems, hereafter called SL3a for the intervening A-rich loop between P2a and P2b stems. A similar secondary structure was reported in early studies of 7SK RNA secondary structure^18,19,21,23^ (**Fig. 2a**). Consistent with the near-equal observed populations, free energy calculations using the RNAeval WebServer^33,34^ showed similar computed free energy (ΔG) values for the two secondary structure states (**Supplementary Table S3**).

The two-state conformational ensemble from DMS-MaPseq data explains the chemical exchange observed in the NMR studies described above. Both SL3e and SL3a states have identical P1 and P3 stems, consistent with the strong NOE connectivities and unambiguous assignments for residues in these stems. The extra observed resonances and ambiguous assignments for resonances of P2 stem residues results from slow exchange of these residues between two distinct secondary structures. DMS-MaPseq shows equal populations of the two states, consistent with similar resonance intensities observed in 2D HSQC spectra. Together, the combined NMR and DMS-MaPseq data show that the SL3 distal end exists as a two-state conformational ensemble.

### SL3 two-state exchange is present in full-length 7SK RNA

To identify if the SL3 two-state conformational ensemble is present in the full-length 332 nt 7SK RNA, we performed DMS-MaPseq experiments using *in vitro* transcribed human 7SK RNA (**Fig. 3a**). Strong agreement was observed for the SL3 distal end between average mutational fraction values of the isolated domain and full-length 7SK RNA constructs (R^2^ = 0.89) (**Supplementary Fig. S4**), indicating that the isolated SL3 domain construct has similar behavior to the full-length 7SK RNA construct. Overall, the DREEM-identified secondary structure of 7SK RNA was nearly identical to previous studies using SHAPE or DMS-based chemical probing experiments, particularly SL1 and SL4^11,18,20,21^ (**Supplementary Fig. S5**). When a single secondary structure is assumed, SL3 has a SL3a-like state, identified by the J1/2 loop and P2a stem, with melted P2b and P3 stems. Similar to the isolated SL3 domain construct, elevated reactivities were observed for P2 stem residues C221, A227, A230, A231, and A251 (**Fig. 3a**). We initially performed DREEM clustering for the DMS-MaPseq data of the entire 332 nt 7SK RNA sequence and found that the secondary structures in the two clusters showed significant differences in the SL1 basal end and SL2 regions (**Supplementary Fig. S5**). The observed populations and secondary structure models are remarkably consistent with previous *in vitro* DANCE-MaP studies of 7SK RNA^11^. The major populated state (71%) resembles the ‘linear’ secondary structure identified in previous studies^18,20,21^. The minor populated state (29%) is similar to the ‘circular’ secondary structure identified in previous studies^11,22,35^, which has an additional SL0 stem from long-range base-pairing between 5’ and 3’ ends.

**Figure 3.**
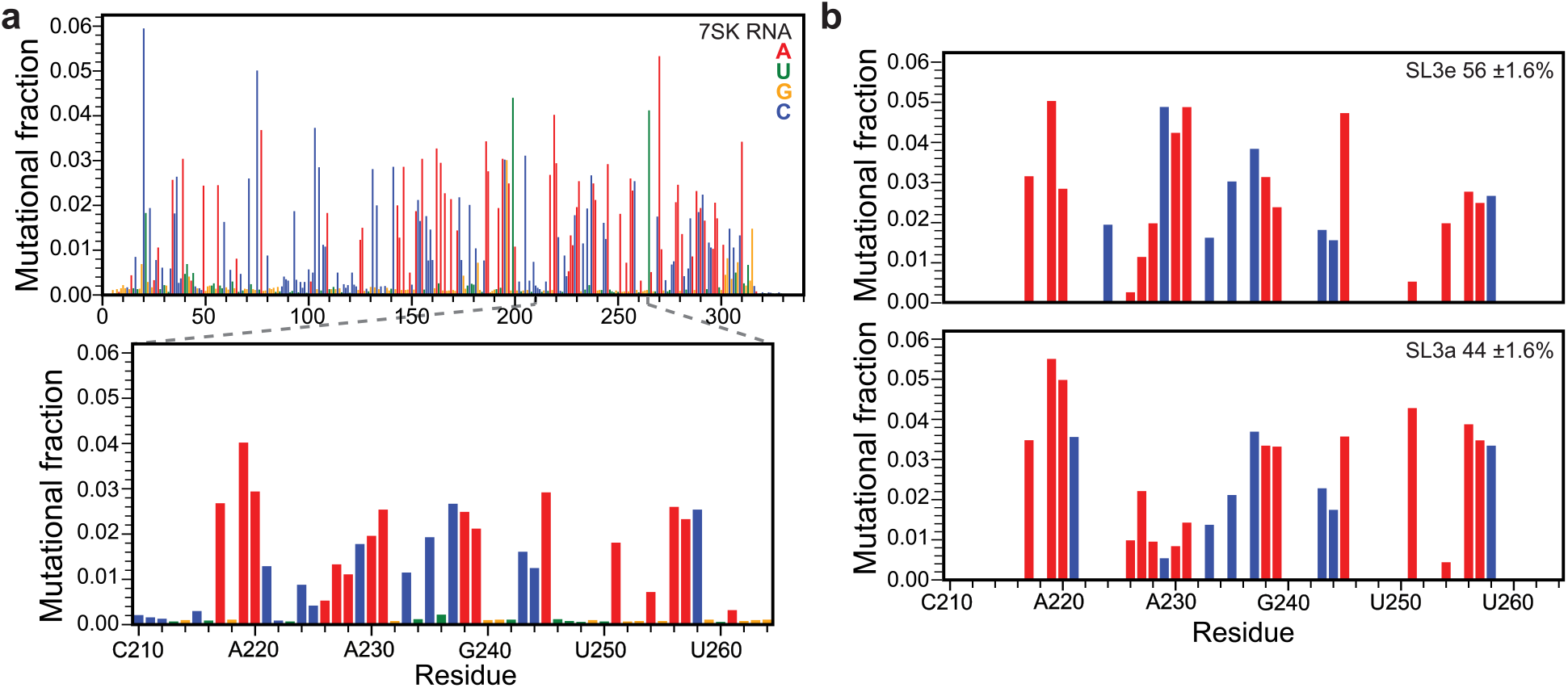
DMS-MaPseq study of *in vitro* full-length 7SK RNA shows two-state conformational exchange. a) *top:* average mutational fraction for full-length 7SK RNA, *bottom:* zoom-in panel of SL3 distal end region corresponding to the isolated domain construct. b) DREEM clusters of SL3a and SL3e conformers are consistent with the domain construct.

Nevertheless, in both clusters, the SL3 distal end shows only the SL3e state. MaP techniques are limited in their ability to deconvolute local structural differences for large RNAs^4^. To address this limitation, we performed DREEM clustering of DMS-MaPseq data from the SL3 distal end (nts 210-264) alone. With this approach, we observed both SL3e and SL3a states with near-equal populations of 54 ± 2% and 46 ± 2%, respectively (**Fig. 3b**, **Table 1, and Supplementary Fig. S4c**). These results demonstrate that the two-state conformational ensemble is present both in the isolated SL3 construct and full-length 7SK RNA.

**Table 1.** SL3 populations determined from DREEM or Linear Combination analysis.

|  | **DREEM** |  | **Linear**<br>**Combination** |  |
| --- | --- | --- | --- | --- |
| **Construct** | SL3e<br>(%) | SL3a<br>(%) | SL3e<br>(%) | SL3a<br>(%) |
| SL3 distal | 52 ± 1 | 48 ± 1 | 54 ± 1 | 46 ± 1 |
| A-lock | 0 ± 0 | 100 ± 0 | na | na |
| E-lock | 100 ± 0 | 0 ± 0 | na | na |
| 7SK RNA | 54 ± 2 | 46 ± 2 | 55 ± 0 | 45 ± 0 |

### Rational construct design locks SL3 into individual states to quench exchange

To further probe the two conformational states and assist in resonance assignment, subdomain constructs were designed for the top and bottom halves of the SL3 distal end (**Supplementary Fig. S7a**). Leveraging 5’-3’ boundary differences between the two states, constructs were designed to stabilize either SL3a (SL3a-top, nts 222-255) or SL3e (SL3e-top, nts 221-253) states (**Fig. 4a**). 1D ^1^H NMR spectra of imino proton resonances showed the expected number of resonances and were able to be assigned by ^1^H-^1^H NOESY experiments (**Fig. 4b and Supplementary** Fig. 6) indicating a single folded state. 2D ^1^H-^13^C HSQC spectra of C2H2 (adenine) resonances showed the expected number of resonances for each construct, further supporting the existence of a single state (**Supplementary Fig. S7b**). Resonances in 2D HSQC spectra of all constructs showed excellent agreement with SL3 distal construct resonances (**Supplementary Fig. S7b**). From NMR spectra of subdomain constructs, resonances in the SL3 distal construct can be assigned to either the SL3e state or SL3a state. For example, the characteristic U223 imino proton resonance at 14.1 ppm that was observed in the SL3 distal construct at elevated temperatures is also observed in SL3a-top, denoted as U223a (**Fig. 4b-c**). In SL3e-top, U223 is shifted upfield to ∼13.6 ppm, denoted as U223e (**Fig. 4c**). SL3e-top showed the characteristic crosspeak between resonances ∼13.8 ppm, assigned as U246e-U247e, observed in SL3 distal at 5 °C in the ^1^H-^1^H NOESY spectrum. Comparison of 2D ^1^H-^15^N HSQC spectra showed that SL3a-top and SL3e-top imino resonances overlay with subsets of SL3 distal resonances, confirming that SL3 distal has slow exchange between two states (**Fig. 4c and Supplemental Fig. S7c**). Three imino resonances were observed in one state but not the other: U246e (SL3e), U250a (SL3a), and G255a (SL3a), and chemical shift perturbations were only observed in the P2 stem (**Fig. 4c-d**). Together, the NMR spectra of the four subdomain constructs superimpose nearly completely onto the SL3 distal construct (**Supplementary Fig. S7**), indicating both SL3a and SL3e states are sufficient to explain the chemical exchange observed in the NMR data of the SL3 distal construct.

**Figure 4.**
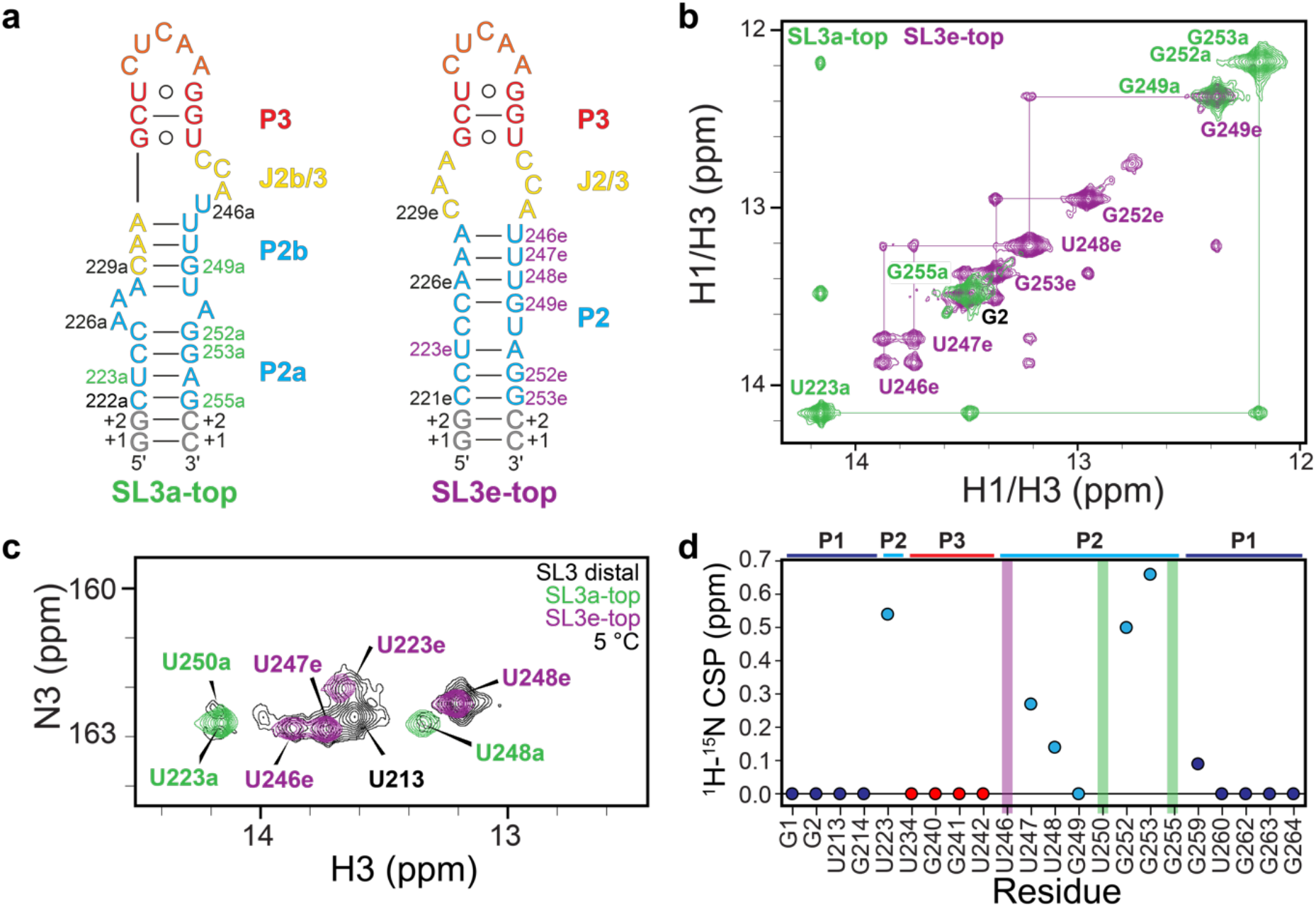
Subdomain constructs of the SL3 top stem lock SL3 into a single state. a) Secondary structures of SL3a-top and SL3e-top constructs. Constructs are colored by secondary structure motifs in SL3e as seen in Figure 2e. b) ^1^H-^1^H NOESY spectrum of imino region of SL3a-top (green) and SL3e-top (purple) constructs. c) ^1^H-^15^N 2D HSQC spectrum of uridine N3H3 resonances. d) Plot of weighted average chemical shift perturbations of SL3e and SL3a imino resonances in SL3 distal construct.

To further characterize each of the SL3 conformational states, we designed point mutations to stabilize either the SL3e or SL3a state (**Fig. 5a-b**). For the locked SL3e construct (E-lock), a C224A substitution replaces the C224•U250 mismatch with a canonical A224-U250 base-pair (**Fig. 5a**). This substitution disfavors the SL3a state by introducing an A•G pair in the SL3a P2a stem. DMS-MaPseq experiments showed 100% SL3e state with high mutational fraction values for reporter residues C229, A230, A231, and A254 (**Fig. 5c**), confirming stabilization of a single state. DMS-MaPseq profiles had excellent agreement between the SL3e cluster in SL3 distal and E-lock constructs (R^2^ = 0.97) (**Fig. 5d**). A comparison of ^1^H-^15^N 2D HSQC spectra of the imino region for SL3 distal and E-lock showed good agreement to SL3e chemical shifts (**Fig. 5e**), with only U223e and U248e, near the mutation site, showing chemical shift perturbations. In addition, a new U250 imino resonance was observed consistent with the new A224-U250 base-pair. Similarly, comparison of ^1^H-^13^C 2D HSQC spectra of adenine C2H2 resonances showed good agreement to SL3e chemical shifts, with a new A224e resonance and chemical shift perturbations to A251e and A226e, near the mutation site (**Fig. 5i**). Importantly, resonances attributed to the SL3a state are not observed in the E-lock construct, consistent with DMS-MaPseq experiments showing a single SL3e state.

**Figure 5.**
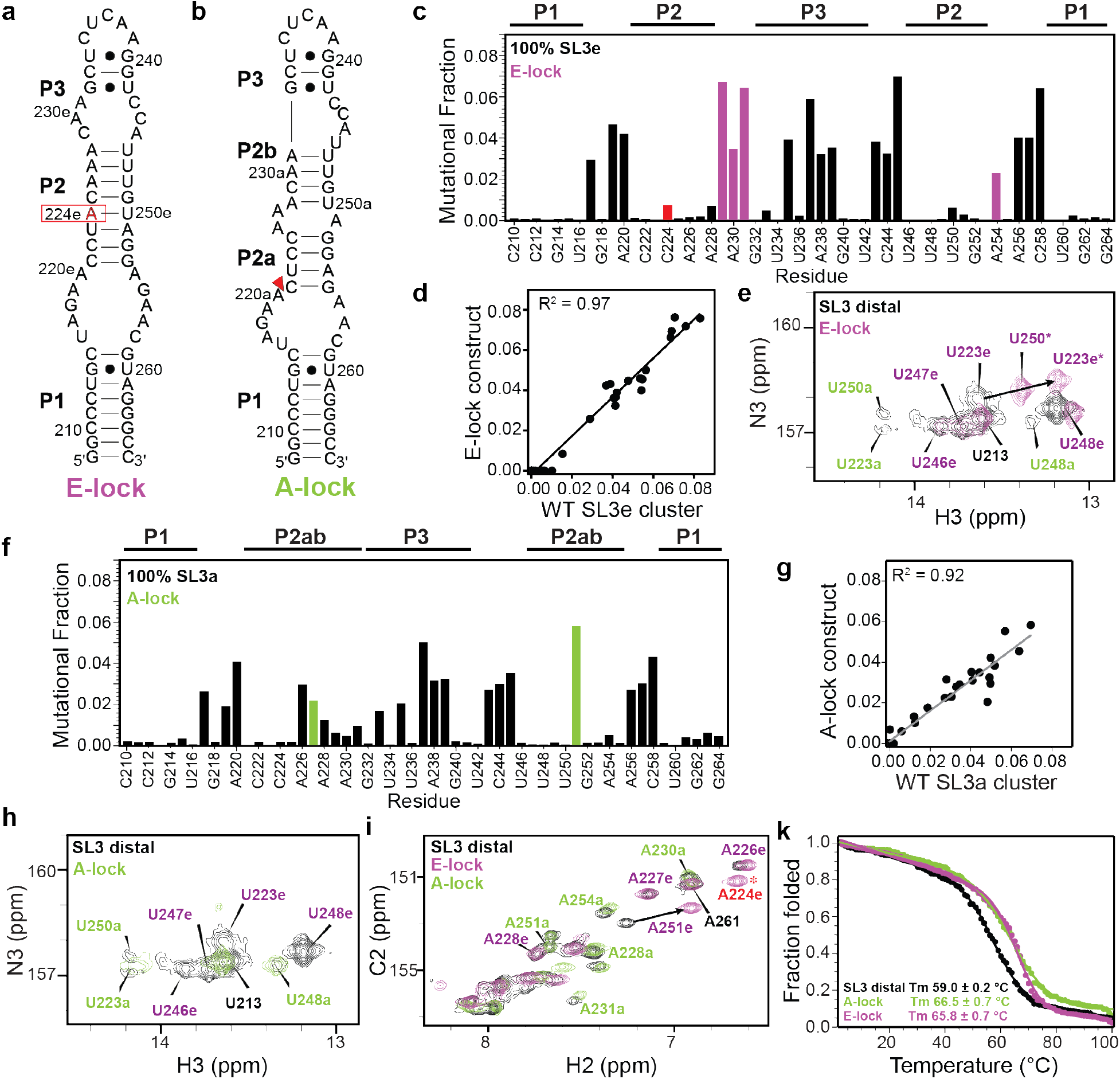
Point substitutions lock SL3 into single conformer. Secondary structure models of SL3 distal constructs a) E-lock and b) A-lock. c) E-lock DMS-MaPseq profile shown as bar plot. Red bar indicates point substitution, purple bars indicate reporter residues identified in Figure 2. DREEM clustering shows 100% SL3e conformer. d) Correlation plot of DMS-MaPseq data of E-lock and SL3 distal construct SL3e cluster show excellent agreement. e) ^1^H-^15^N 2D HSQC of uridine imino resonances of SL3 distal (black) and E-lock (purple) show good agreement with superimposition of resonances corresponding to SL3e conformer. f) DMS-MaPseq profile of A-lock shown as bar plot. Green bars indicate reporter residues identified in Figure 2. DREEM clustering shows 100% SL3a conformer. g) Correlation plot of DMS-MaPseq data of A-lock and SL3 distal construct SL3a cluster show excellent agreement. Open circle indicates outlier residue A219. h) ^1^H-^15^N 2D HSQC of uridine imino resonances of SL3 distal (black) and A-lock (green) show good agreement with superimposition of resonances corresponding to SL3a conformer. i) ^1^H-^13^C 2D HSQC of adenine C2H2 resonances of SL3 distal (black), E-lock (purple), and A-lock (green). j) Thermal unfolding profiles of SL3 distal (black), E-lock (purple), and A-lock (green).

To stabilize the SL3a state, we performed a deletion of C221 (A-lock) to the SL3 distal construct (**Fig. 5b**). C221 is located in the J1/2 loop in SL3a and is the closing base-pair in SL3e. We predicted that this loop deletion would minimally impact the SL3a state and disfavor the SL3e state by eliminating one base-pair. DMS-MaPseq experiments showed 100% SL3a state with high mutational fraction values for A227 and A251 (**Fig. 5f**) and excellent agreement to the SL3a cluster in the SL3 distal construct (R^2^ = 0.92) (**Fig. 5g**). The ^1^H-^15^N 2D HSQC spectrum of the imino region showed excellent agreement to SL3a chemical shifts (**Fig. 5h**), in support of formation of a single SL3a state. Inspection of ^1^H-^13^C 2D HSQC spectra of adenine C2H2 resonances shows excellent agreement to SL3a resonances (**Fig. 5i and Supplementary Fig. S9**) with minimal chemical shift perturbations. Resonances assigned to SL3e are not observed in the A-lock construct, consistent with DMS-MaPseq data showing a single SL3a state. Together, the subdomain and point mutant constructs in this study validate the presence of two distinct states in the SL3 P2 stem and demonstrate that SL3 can be locked into a single conformer with minimal structural perturbations to either conformer.

### Locking SL3 distal into a single state increases thermal stability

To evaluate the thermodynamic stability of SL3 distal constructs, optical melting was performed using circular dichroism (CD) spectroscopy. A single cooperative melting transition was observed with a melting temperature (Tm) of 59.0 ± 0.2 °C for the wild-type SL3 distal construct (**Fig. 4j and Supplementary Fig. S10**). Surprisingly, both E-lock and A-lock constructs showed increased melting temperatures with Tm values of 66.5 ± 0.7 °C and 65.8 ± 0.7 °C, respectively. The similar Tm values for the locked constructs are consistent with SL3a and SL3e states having similar free energies.

### Both SL3e and SL3a states are conserved among vertebrates

SL3 is a conserved structural domain identified in vertebrates, hexapoda, and coleoptera ^22,36,37^. For the human 7SK RNA SL3 distal end, we found that the RNAeval webserver^33^ computed a free energy difference (ΔΔG) between SL3e and SL3a states to be approximately ∼0.4 kcal/mol, similar to the estimated ΔΔG assuming Boltzmann statistics, suggesting that the thermodynamic parameters used in the ViennaFold package perform reasonably well at predicting the energy differences between the two secondary structures.

To investigate whether the SL3 two-state conformational ensemble is specific to humans, or conserved more broadly across vertebrates, we used previously identified vertebrate 7SK RNA sequences from Gruber et al^37^ and RNAsubopt^33^ to predict the five lowest-energy secondary structures for the SL3 distal end sequence. Consistent with prior studies, we found high sequence conservation among 7SK RNA sequences^22,37^ that is compatible with both SL3a and SL3e states (**Fig. 7a, Supplementary Fig. S13b**). In particular, all mammals except for rodents and marsupials had identical sequences to human and, as expected, both SL3a and SL3e states were predicted with similar ΔΔG values (**Supplementary Table S3, Fig. S14)**. In rodents, U246 is absent, resulting in the SL3a state having significantly lower energy compared to the SL3e state (**Supplementary Fig. S13b, S14**). Consistent with this finding, the SL3a secondary structure was reported in prior chemical probing studies of 7SK RNA in rat Novikoff hepatoma cells ^23^ and mouse embryonic stem cells ^19^. In marsupials, an 8-nt deletion of P2 stem residues C222-C229 results in a unique secondary structure that is neither SL3a nor SL3e (**Supplementary Fig. S14**). Across fishes, with the exception of *Danio rerio* (zebrafish), both states were predicted with equal energies (**Supplementary Fig. S13b, S14**). For *Danio rerio*, which has insertions in J1/2 and J2/3 loops, only the SL3e state is predicted to form. Compared to human, *Mustelus* and *Lampetra* have substantial sequence differences and lack J1/2, J2/3, and apical loop sequences despite having respective SL3e-like and SL3a-like features in the P2 stem region. Taken together with our mutagenesis studies, the phylogenetic analysis shows remarkable sensitivity in the sequence dependence on the SL3 secondary structure. The high degree of evolutionary conservation of both SL3e and SL3a states in 7SK RNA SL3 suggests a functional role for both states.

### A two-state physical model is sufficient to explain NMR and chemical probing data

At present, MaP-based approaches are limited by a reliance on correlated mutations and thermodynamics-guided secondary structure prediction^4,38^. We sought to analyze DMS-MaPseq data independently of DREEM to provide additional validation for the observed two-state conformational ensemble. Given the excellent fit in DMS-MaPseq profiles between SL3 distal DREEM clusters and locked constructs (**Fig. 5d, g**), we reasoned that these mutants could be used as ‘fingerprints’ to represent either the SL3a or SL3e states. To evaluate whether the DMS-MaPseq data could be fit to a two-state model, we generated in-house software that computes populations of two states using linear combination. Here, the DMS-MaPseq profiles of the A-lock and E-lock constructs were used as templates for SL3a and SL3e conformers, respectively. For both SL3 distal (**Fig. 6a**) and 7SK RNA (**Fig. 6b**) constructs, populations fit to ∼55% for SL3e and ∼45% for SL3a consistent with DREEM calculations (**Table 1**). The correlation between experimental and fit data was excellent, with R^2^ = 0.99 for the isolated SL3 domain and R^2^ = 0.87 for SL3 in full-length 7SK RNA. These results show that the DMS MaPseq data can be explained by a physical two-state model in which SL3 exists as two distinct states, and validates the populations determined using DREEM.

**Figure 6.**
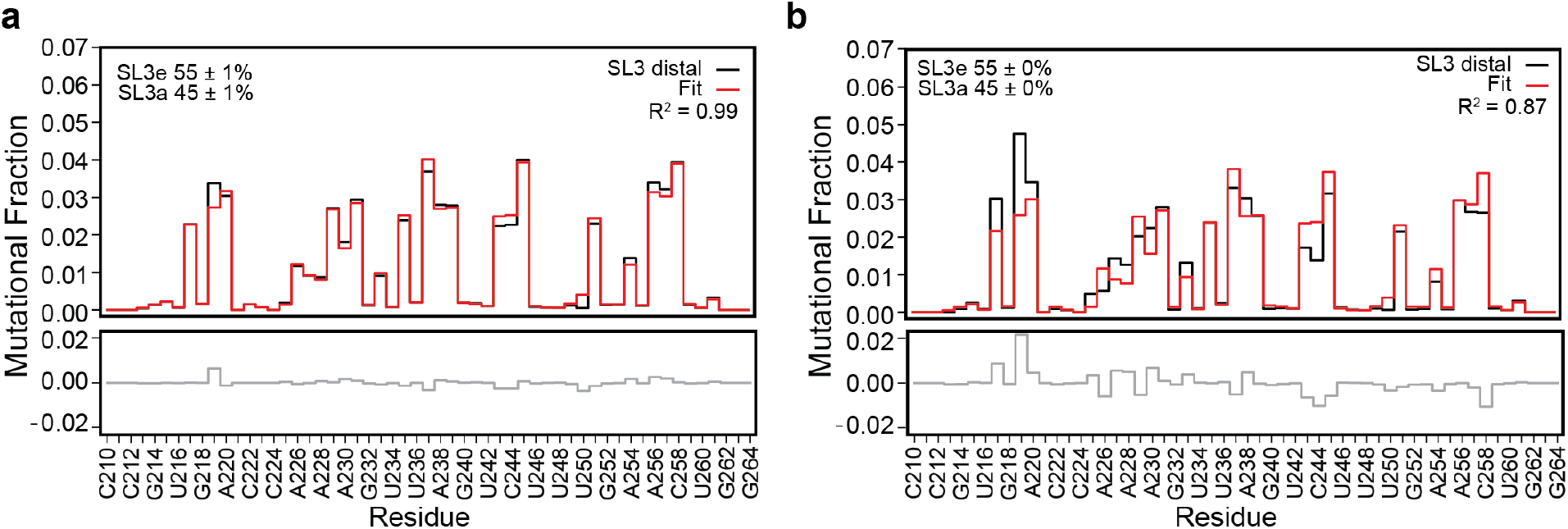
Linear combination analysis fits a two-state physical model. a) SL3 distal construct and b) SL3 region of full-length 7SK RNA. DMS-MaPseq mutational fraction values colored black, fit line colored red, and residual colored gray.

## Discussion

RNA conformational exchange and secondary structural rearrangements play essential roles in protein recruitment and cellular function^1,39,40^. Despite this significance, there are currently limited methods capable of measuring these structural dynamics. The SL3 domain of 7SK RNA was previously identified to be a conserved structural element of 7SK RNA^36,37^, with prior studies reporting either SL3e^11,20^ or SL3a^19,21,23^ states. Here using combined NMR, DMS-MaPseq, optical melting, and mutagenesis approaches, we show that both states are populated in human 7SK SL3 RNA as a two-state conformational ensemble. A previous NMR study of the isolated SL3 domain reported the SL3e state^20^; however, differences in NMR field strength and buffer conditions may account for the reported differences. Prior *in vitro* and *in cellulo* chemical probing studies that identified the SL3e state showed elevated chemical reactivity in P2 stem residues^11,20^, particularly residues C221 and A251 which we identify as reporter residues for the SL3a state. Our data shows that this reactivity is due to conformational exchange between SL3e and SL3a states. Our study reconciles disparities in the literature by showing that SL3 exists as a two-state conformational ensemble in human 7SK RNA, explaining why in some studies the SL3e state is identified and the SL3a state is identified in others. In addition, our phylogenetic analysis provides an explanation for mouse and rat studies that report the SL3a state.

While rearrangements of base-pairs within stems have been observed^41–43^ there is often a substantial difference in free energy resulting in a major populated state and minor alternative state. In sharp contrast, SL3 has similar free energies in the two states and near-equal populations. Likewise, bistable RNAs with large-scale switching between secondary structures have been observed in rationally designed RNAs^44,45^ and riboswitch expression platforms^46^. For these systems, switching typically takes place via strand exchange between an RNA hairpin and single-stranded overhang, not within a stem of a hairpin. The transition between the two states in the SL3 distal end involves a rearrangement of eight base-pairs and is predominantly localized in the P2 stem (**Fig. 7b**). Evolutionary conservation of SL3e and SL3a secondary structures suggests a potential functional role of both states. The two states significantly differ in loop sequence and composition: while SL3e has symmetric J1/2 and J2/3 loops, SL3a has asymmetric J1/2 and J2/3 loops and a new A-rich loop in the P2ab stem (**Figs. 2e, 7b**). These differences may be functionally relevant, as loops often serve as protein recognition sites. 7SK SL3 is required for P-TEFb release from 7SK RNP ^26,27^ and several proteins have been identified to bind SL3 (e.g. hnRNPA1, SRSF2, RBM7) in the P-TEFb release pathway^20,24,25,47^.

**Figure 7.**
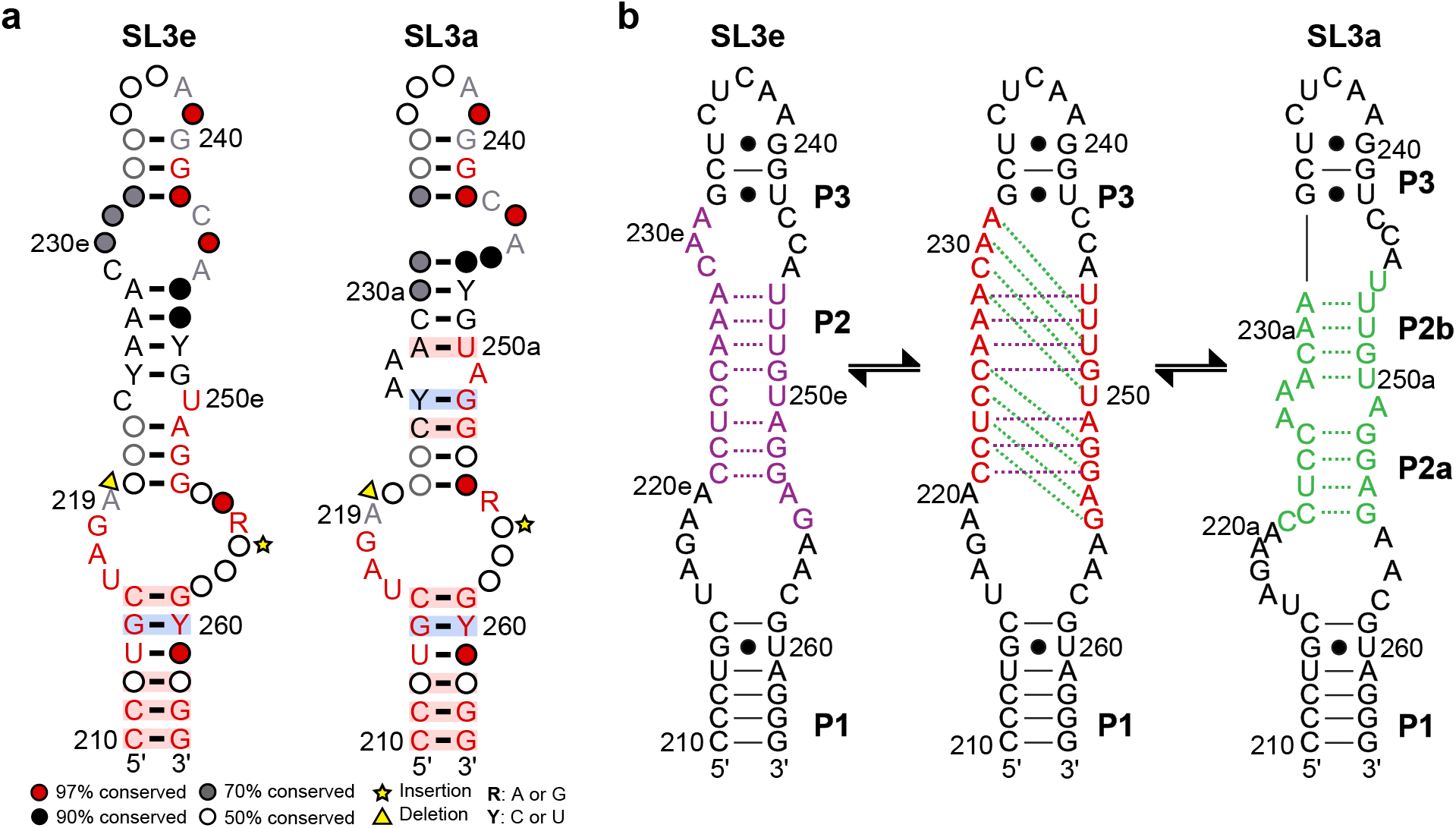
7SK SL3 sequence conservation and exchange model. a) Sequence conservation plotted on SL3e and SL3a states using R2R^69^ shows conservation of both potential states. b) Model of exchange between SL3e and SL3a states. Residues that are involved in base-pair rearrangements are shown in purple (SL3e), red (intermediate), and green (SL3a).

In summary, this study highlights the power of combined application of NMR and chemical probing to provide quantitative insights into RNA conformational ensembles. We demonstrate that local base-pair rearrangements may be missed in current chemical probing approaches of large RNAs and anticipate that other functionally relevant RNAs may undergo similar structural transitions.

## Methods

### Sample preparation

Chemically synthesized DNA templates of SL3 domain constructs were purchased from Integrated DNA Technologies (IDT) with a 2’ O-methyl modification for the two nucleotides at the 5’ end. The DNA template for full-length 7SK RNA sequence (NCBI Gene ID 125050), containing a T7 RNAP promoter sequence, was cloned into a pUC19 plasmid. Linear DNA template was obtained from PCR amplification of the DNA template region and purified using the DNA clean and concentrator spin-column kit (Zymo). RNA samples were prepared by *in vitro* transcription (IVT) using T7 RNA polymerase (Addgene #124138^48^, prepared in-house). T7 RNA Polymerase, 500 nM DNA template, and transcription buffer (40 mM Tris pH 8, 1 mM spermidine, 0.01% Triton-X, 40 mM MgCl_2_, 2.5 mM DTT, 20% DMSO, and 2 mM each rATP, rCTP, rUTP, rGTP) were incubated at 37 °C for 6-8 hours. Distal RNA constructs were purified by 15% denaturing polyacrylamide gel electrophoresis (PAGE) and full-length 7SK RNA constructs were purified by 5% denaturing PAGE. The RNA band was visualized by UV shadowing with a handheld UV lamp at 254 nm. After band excision, RNA was eluted from the gel using the ‘crush and soak’ method ^49^ by incubating gel pieces in crush and soak buffer (300 mM sodium acetate pH 5.2, 1 mM EDTA) for 24-48 hours at room temperature. RNA was further purified to remove acrylamide contaminants by ion-exchange chromatography using a diethylaminoethanol (DEAE) column (GE Healthcare) and elution into buffer (10 mM sodium phosphate pH 7.6, 1 mM EDTA, 1.5 M KCl). RNA was diluted to <100 μM in ultrapure water and annealed by heating to 95 °C for 3 minutes, followed by snap cooling on ice for 1 hour. RNA was then buffer exchanged into the appropriate buffer using a 3-10 kDa Amicon concentrator (Millipore Sigma).

SL3 distal constructs for chemical probing were prepared using established methods^50^. Constructs were designed with flanking GGAAGATCGAGTAGATCAAA and AAAGAAACAACAACAACAAC sequences at the 5’ and 3’ ends, respectively, for primer annealing during downstream PCR of dsDNA and reverse transcription of cDNA (**Table S2**). Primers for PCR assembly of the DNA templates were designed using Primerize^51^ (https://primerize.stanford.edu). PCR was performed using Q5 DNA Polymerase (NEB) followed by purification using DNA clean and concentrator spin-column purification (Zymo). RNA was transcribed using IVT as described above, purified using gel excision, and stored at -80 °C in ultrapure water until use.

### NMR spectroscopy

Solution NMR spectroscopy experiments were performed at 278.15 K and 298.15 K on a Bruker Neo 600 MHz NMR spectrometer equipped with a triple-resonance HCN cryoprobe. NMR samples were prepared in NMR buffer (20 mM sodium phosphate, 50 mM KCl, pH 6.0) with added 5% D_2_O at 0.1-0.8 mM concentrations in 3 mm NMR tubes (Norell). Exchangeable (H1, H3, H41, H42) and nonexchangeable (H2, H5, H6, H8, H1’) proton resonances were assigned using 2D ^1^H-^1^H NOESY spectra of unlabeled RNA samples with mixing times of 150 ms, 200 ms, and 250 ms. ^1^H-^15^N HSQC and ^1^H-^13^C HSQC spectra were collected using ^13^C/^15^N labeled RNA samples. Data were processed using NMRPipe^52^ and analyzed using NMRFAM-Sparky 1.470 powered by Sparky 3.190^53^ in the NMRbox virtual machine^54^. Weighted average chemical shift perturbations (CSP) were calculated using the equation 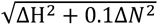 ^55^.

### Circular Dichroism (CD) spectroscopy

CD experiments were performed on a Jasco815 spectrometer equipped with a Peltier temperature control device. RNA samples were prepared at 25 μM in CD buffer (20 mM sodium phosphate, 50 mM KCl, 0.1 mM EDTA, pH 6.0). Thermal unfolding experiments were performed in triplicate with a temperature range of 2 °C – 100 °C (forward) or 100 °C – 5 °C (reverse) and a ramp rate of 1 °C/min, 2 nm bandwidth, and molar ellipticity values measured at 258 nm every 1 °C. Data was fitted and visualized using an in-house Python script adapted from the Delta-melt program developed by Al-Hashimi and coworkers^56,57^. Data were fit assuming a two-state folding model^58^ where the mean ellipticity is:

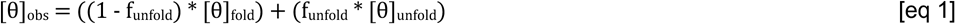

Assuming the mean ellipticity value to be a linear function of temperature, this equation can be expanded into:

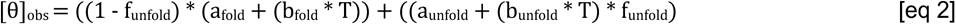

Where:

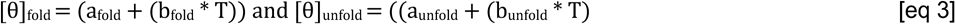

a_fold_, b_fold_, a_unfold_ and b_unfold_ are linear coefficients that represent the slope and intercept of the RNA folded or unfolded states as a function of temperature, T is the temperature (K), and f_unfold_ is the fraction of the RNA folded/unfolded. f_unfold_ can be determined using the equation:

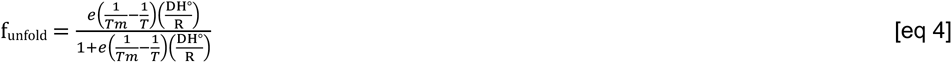

Where Tm is the melting temperature (K), ΔH° is the enthalpy, and R is the gas constant.

### DMS-MaPseq

DMS-MaPseq experiments were performed using established protocols^50^. RNA samples were annealed by heating at 90 °C in water for two minutes then 4 °C for five minutes, followed by addition of 1x folding buffer (100 mM HEPES, pH 8.0) at 25 °C. A 15% DMS (Sigma Aldrich) solution in ethanol was used for DMS-MaPseq experiments. 7.5 pmoles RNA was incubated with 0.75-1.5% DMS solution at 25 °C for six minutes, then quenched with β-mercaptoethanol (BME). As a control, reactions were performed using an equivalent volume of ethanol to estimate mutations induced by reverse transcriptase enzyme in the absence of DMS. DMS-treated samples were purified using RNA clean and concentrator-5 spin-column kits (Zymo). Complementary DNA (cDNA) was generated using either TGIRT-III (In-Gex) or Marathon (Kerafast) reverse transcriptases. RNA was reverse transcribed using 100 mM DTT, 10 mM dNTPs, RT buffer (250 mM Tri-HCl pH 8.0, 375 mM KCl, 15 mM MgCl_2_), primers with custom barcodes for downstream demultiplexing (**Table S2**) and TGIRT-III or Marathon RT enzyme. For TGIRT-III, the reaction was incubated at 57 °C for two hours. For Marathon, the reaction was incubated at 42 °C for three hours. The reaction was quenched with NaOH, heated at 90 °C for 2 minutes then cooled at 4 °C, followed by the addition of acid to neutralize the reaction (5 M NaCl, 2 M HCl, 3 M NaAc). The cDNA was purified using Zymo oligo clean and concentrator-5 kit (Zymo). To prepare dsDNA for sequencing, purified cDNA was used as a template for PCR using Q5 DNA Polymerase (NEB) and forward and reverse sequencing primers (**Table S2**). For SL3 distal end constructs, PCR was performed using 16 cycles with a 15 second extension time. For full-length 7SK RNA, PCR was performed using 20 cycles with a 20 second extension time. An annealing temperature of 62 °C was used for both SL3 distal end constructs and full-length 7SK RNA. PCR products were visualized on a 4% agarose E-gel (Invitrogen) and gel purified using an Invitrogen gel purification kit (Invitrogen) following the manufacturer’s protocol. The dsDNA samples were quantified using a qubit 1x dsDNA high-sensitivity assay kit (Invitrogen) and qubit flex fluorometer (Invitrogen). Library preparation was performed by diluting the dsDNA samples to ∼1 nM and mixing individual samples of interest for up to 395 pmol of dsDNA. For the final library, 39 pmol of dsDNA was mixed with 36 pmol PhiX (Illumina) for sequencing. SL3 distal, E-lock, and A-lock samples were sequenced on the iSeq 100 (Illumina) using iSeq 100 i1 reagent v2 kit (300 cycle) 2 x 150 read length (Illumina). The total number of reads were 150,000-400,000 per sample. Full-length 7SK RNA was sequenced on the miSeq (Illumina) at the UNMC Sequencing core using miSeq reagent kit v3 (600 cycles) 2 x 300 read length (Illumina). The total number of reads were 1-2 million reads per sample.

### Statistical analysis and visualization

Data analysis for DMS-MaPseq was performed using DREEM as previously described in Tomezsko *et al.*^32^. To account for the stochastic nature of the DREEM algorithm, we performed three iterations of DREEM clustering for each experimental replicate to calculate the average population and standard deviation. Data was visualized using in-house python scripts (https://github.com/ceichhorn2) that utilize SciPy^59^, Pandas^60^, Matplotlib^61^, NumPy^62^, Seaborn^63^ and Biopython^64^ module. Final secondary structure models were prepared using Varna^65^.

### Linear combination analysis

Linear combination analysis was performed using the DMS mutational fractions of the E-lock and A-lock constructs as ‘fingerprints’ to generate a reference dataset. To account for end fraying, the terminal three residues at the 5’ and 3’ ends were set to zero. Additionally, mutation sites in the E-lock (C224) and A-lock (C221) constructs were set to zero for SL3 distal or 7SK RNA (nts 210-264), E-lock, and A-lock mutational fraction data. To account for variations in DMS treatment, a weighting factor was applied to scale the magnitude of the reference dataset to the SL3 distal or 7SK RNA (nts 210-264) data. The resulting calculation determines the populations of the SL3e and SL3a states for the input DMS-MaPseq data from a linear combination of the reference datasets fit to the experimental data. The residual of the fit was determined by taking a difference between the experimental data and linear combined fit values.

### Phylogenetic analysis of 7SK RNA SL3

Multiple sequence alignment (MSA) for 7SK RNA SL3 vertebrate sequences (**Table S3**) from Gruber et al^37^ was performed using LocARNA v1.9.1 (https://github.com/s-will/LocARNA)^66,67^. Suboptimal 7SK RNA SL3 secondary structure prediction for each vertebrate organism was performed using RNAsubopt 2.6.2^33^. Free energies (ΔG) of the SL3a and SL3e states in the top five predicted secondary structures were computed at 25 °C using Andronescu et al 2007 RNA parameter model^68^ using RNAeval^33^. The sequence conservation from the MSA was drawn on SL3e and SL3a states using R2R (https://sourceforge.net/projects/weinberg-r2r/)^69^. The dot bracket secondary structure notation in the stockholm file was modified to either SL3e or SL3a secondary structures for R2R. Due to large gaps in *Lampetra* and *Petromyzon*, the line ‘#=GF R2R keep allpairs’ was added to the stockholm file to prevent errors from the secondary structure drawing.

## Data availability

Python scripts are deposited in GitHub (https://github.com/ceichhorn2). Processed DMS-MaPseq data is deposited in the RNA Mapping Database (RMDB) under RMDB_ID ####. Raw demultiplexed sequencing data is deposited in the NCBI Sequence Read Archive (SRA) under Accession ####.

## Acknowledgements

We gratefully acknowledge Addgene for the pQE30-His-T7RNAP plasmid used to prepare T7 RNA Polymerase enzyme for *in vitro* transcribed RNA samples, which was a gift from Sebastian Maerkl & Takuya Ued (Addgene plasmid #124138). We thank Dr. Robert Peterson and Dr. Martha Morton for NMR assistance. We thank Dr. Jennifer Bushing and the University of Nebraska Medical Center Genomics Core.

## Author contributions

CDE conceived and oversaw all aspects of the project. MBC, JDY, and CDE designed experiments and performed data analysis. MBC prepared samples and performed experiments. BL performed library preparation for DMS-MaPseq experiments. JDY wrote the linear combination analysis software. MBC and CDE wrote the paper with input from all authors.

---

**Figure S1.**
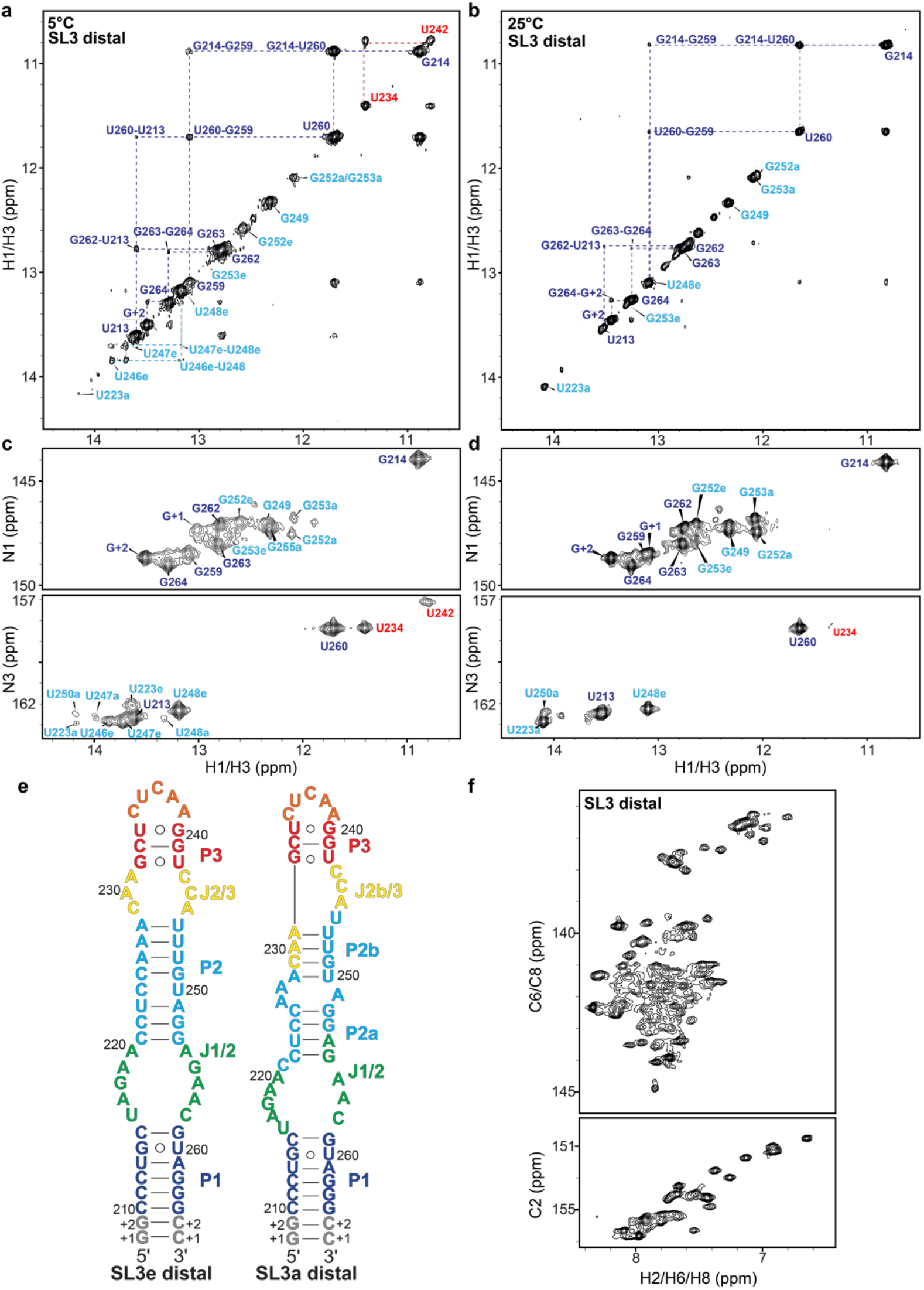
NMR spectra of SL3 distal construct. ^1^H-^1^H NOESY spectrum of imino proton resonances at a) 5 °C or b) 25 °C shows connectivities for P1 stem resonances. ^1^H-^15^N HSQC spectra at c) 5 °C or d) 25 °C shows resonances for both SL3e and SL3e states. P3 stem residues are observed at 5 °C, colored red. e) Secondary structures of SL3e or SL3a states. f) ^1^H-^13^C HSQC spectrum of nucleobase (C2H2, C6H6, C8H8) resonances show greater than the expected number of resonances.

**Figure S2.**
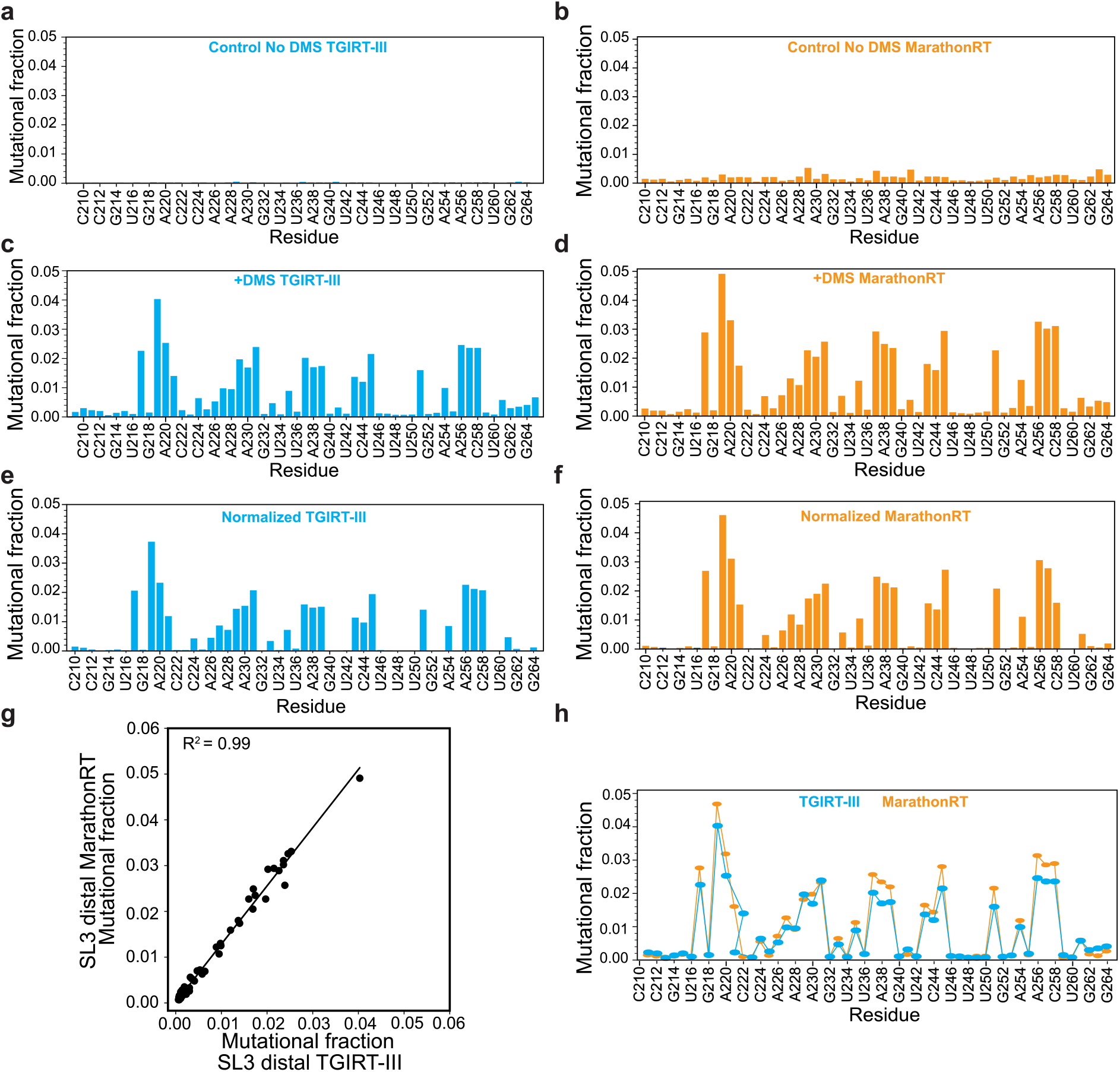
Comparison of TGIRT-III and Marathon RT activity in DMS-MaPseq experiments for SL3 distal constructs. a) and b) control DMS-MaPseq for SL3 distal without DMS using TGIRT-III or MarthonRT for reverse transcription. c) and d) DMS-MaPseq for SL3 distal with DMS using TGIRT-III or MarathonRT. e) and f) Baseline subtracted (normalized) mutational fraction profiles for TGIRT-III or MarathonRT. g) Correlation plot between the normalized TGIRT-III and normalized MarathonRT data. h) Overlay of normalized TGIRT-III and normalized MarathonRT mutational fraction.

**Figure S3.**
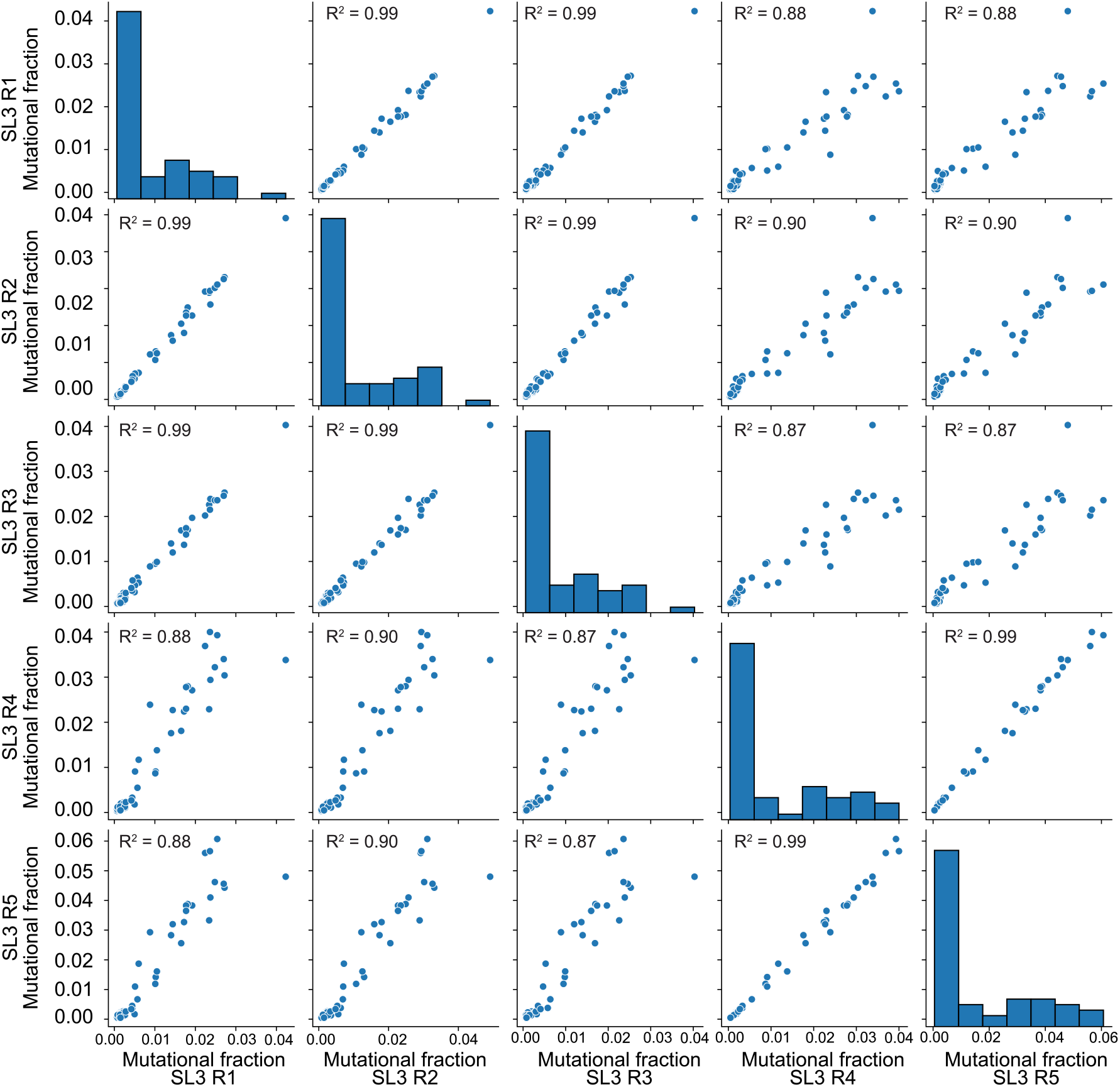
Comparison of DMS-MaPseq data for five independent replicates of SL3 distal constructs shows a strong correlation among all replicates.

**Figure S4.**
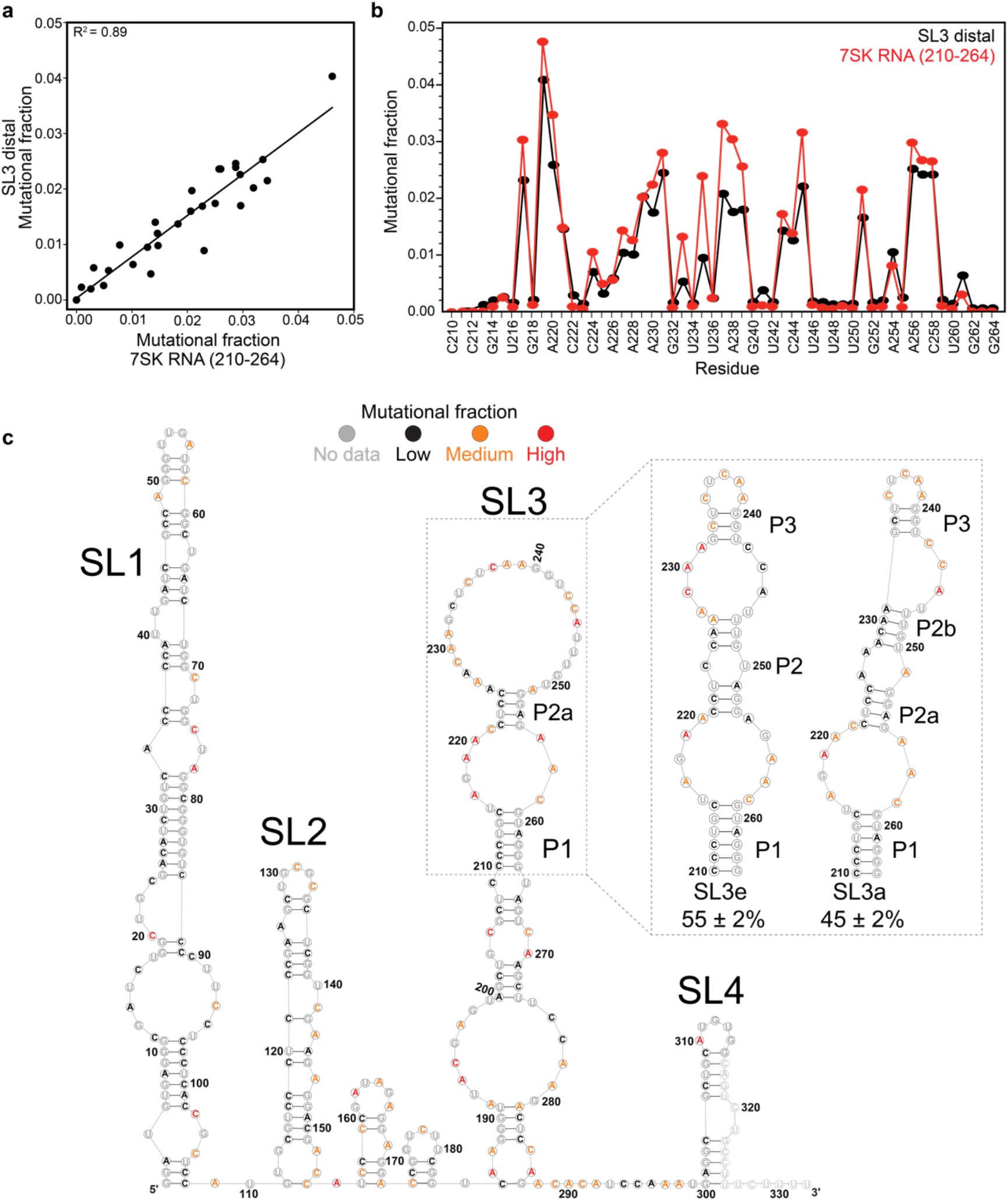
Comparison of SL3 domain in isolated SL3 construct and full-length 7SK RNA, and DREEM analysis of 7SK RNA (nts 210-264). a) DMS-MaPseq data for the isolated SL3 construct and full-length 7SK RNA plotted a) as a correlation plot and b) per-residue with SL3 distal domain construct colored black and corresponding residues in the full-length 7SK RNA colored red. c) Secondary structure of 7SK RNA from DREEM using mutational fraction of 7SK replicate 1 as a representative example. Low, medium and high mutational fraction thresholds were determined from DREEM. *Inset:* DREEM clustering analysis using the 210-264 nt region of the 7SK RNA shows SL3e and SL3a states.

**Figure S5:**
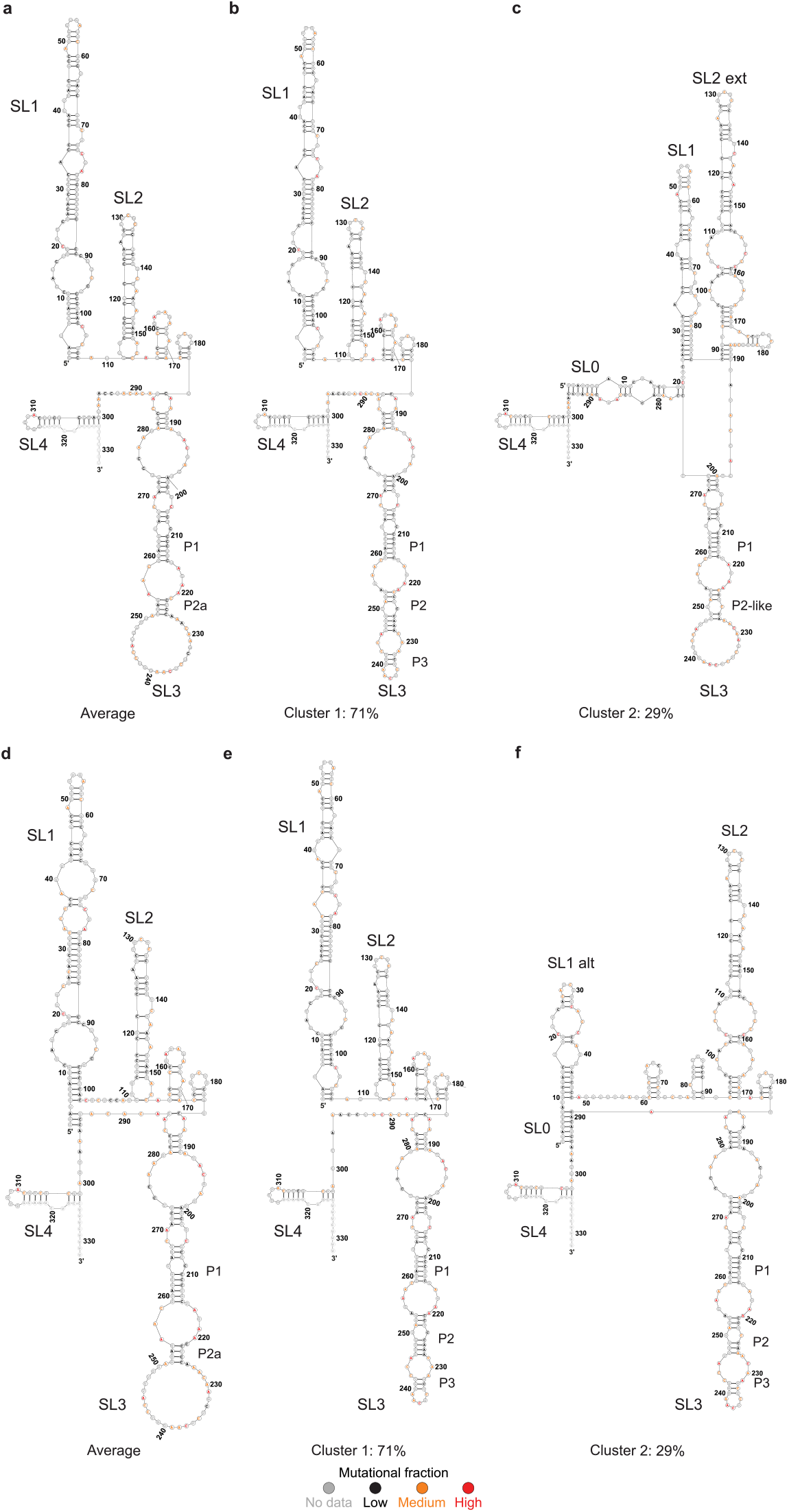
DREEM clustering analysis of full-length 7SK RNA (1-332 nt). a) and d) DREEM-identified secondary structure of full-length 7SK RNA replicate 1 and replicate 2 before clustering, respectively. b), c) and e), f) Secondary structure of full-length 7SK RNA after clustering into two states using DREEM. Grey, black, orange or red denotes; no data, low, medium or high DMS mutational fraction thresholds determined from DREEM.

**Figure S6.**
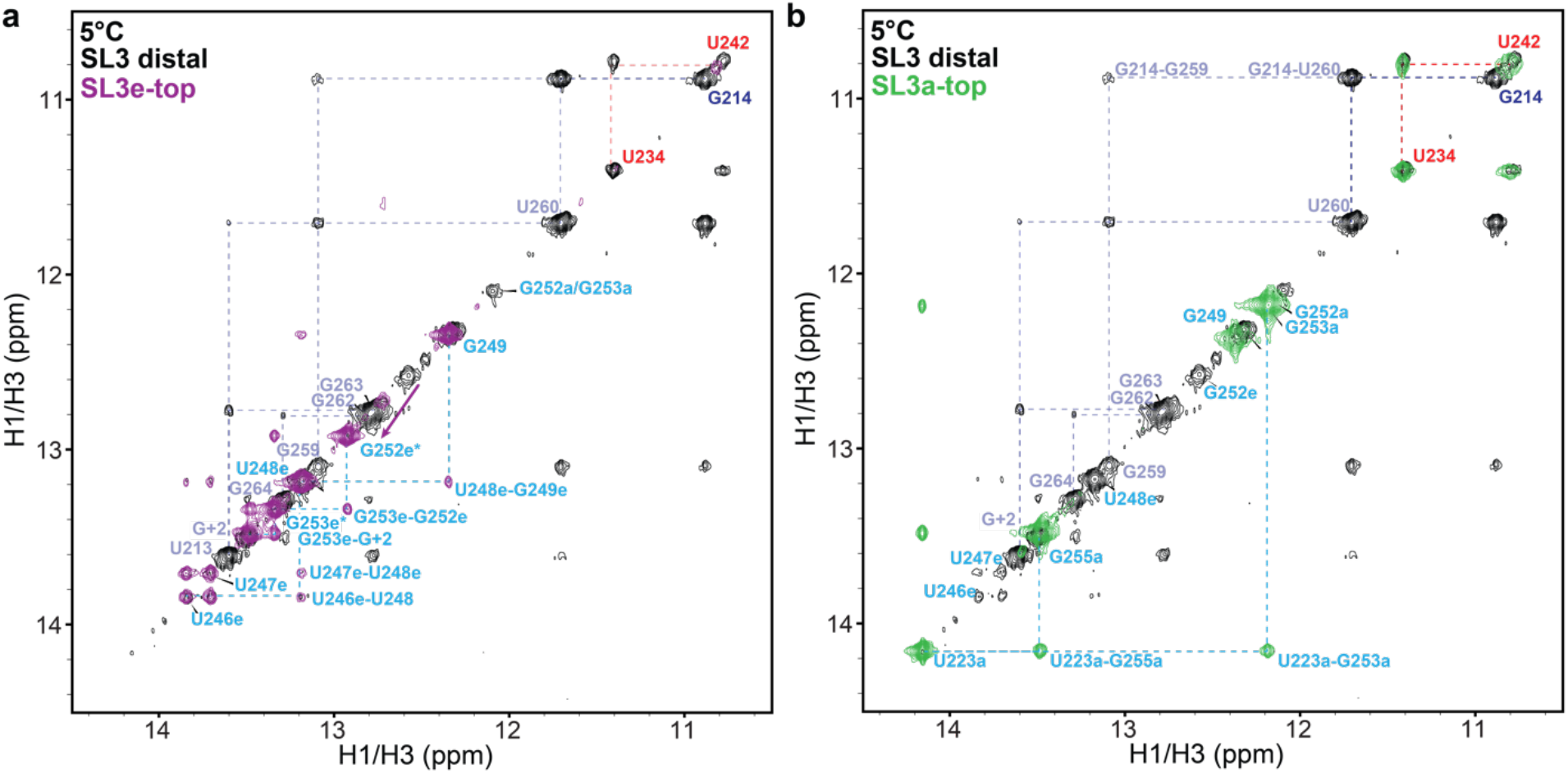
Imino proton resonance assignments for SL3a-top and SL3e-top constructs. a) ^1^ H-^1^H NOESY spectrum of SL3e-top (purple) overlaid with SL3 distal (black). b) ^1^ H-^1^H NOESY spectrum of SL3a-top (green) overlaid with SL3 distal (black).

**Figure S7.**
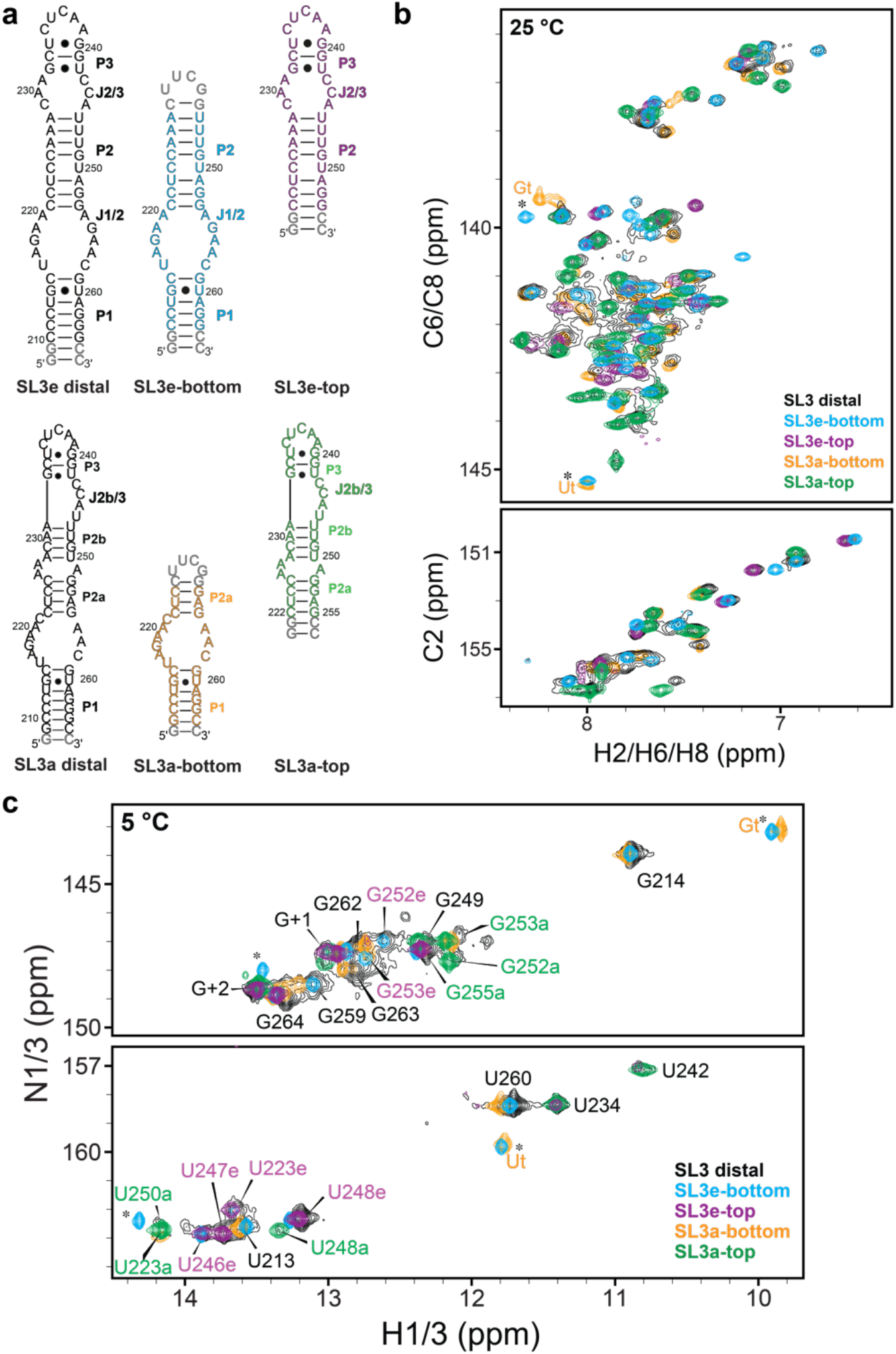
2D HSQC spectra for SL3 subdomain constructs. SL3e or SL3a bottom constructs are capped with cUUCGg tetraloop. Resonances for non-native cUUCGg tetraloop for SL3e- and SL3a-bottom constructs are indicated with asterisks. a) Secondary structures of SL3 domain constructs. b) ^1^H-^13^C HSQC spectrum of aromatic (C2H2, C6H6, C8H8) resonances, shows that together subdomain constructs superimpose onto SL3 distal constructs. c) ^1^H-^15^N HSQC spectrum of imino resonances shows near-complete superimposition of subdomain constructs onto SL3 distal constructs. Residue labels indicate SL3e state (purple) or SL3a state (green) resonances. P1 and P3 stem residues are labeled in black.

**Figure S8.**
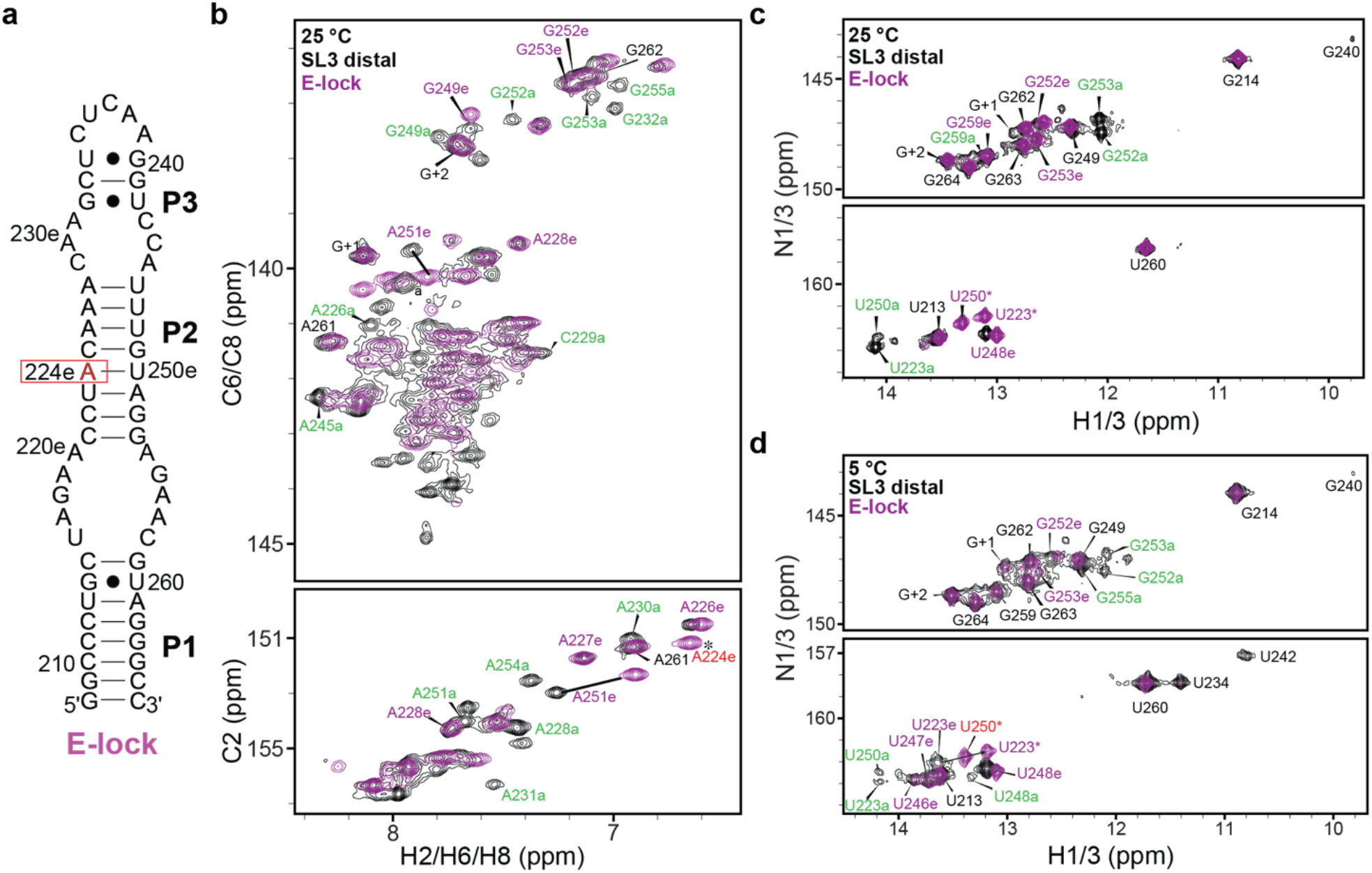
NMR data of SL3 E-lock construct. a) Secondary structure of SL3 E-lock construct with mutation site indicated with a red box. b) 2D ^1^H-^13^C HSQC spectra of SL3 E-lock construct (purple) overlaid on wild-type SL3 distal construct (black). c-d) Overlay of 2D ^1^H-^15^N HSQC spectra of SL3 E-lock and wild-type SL3 distal construct. Resonance labels for SL3e state (purple) and SL3a state (green).

**Figure S9.**
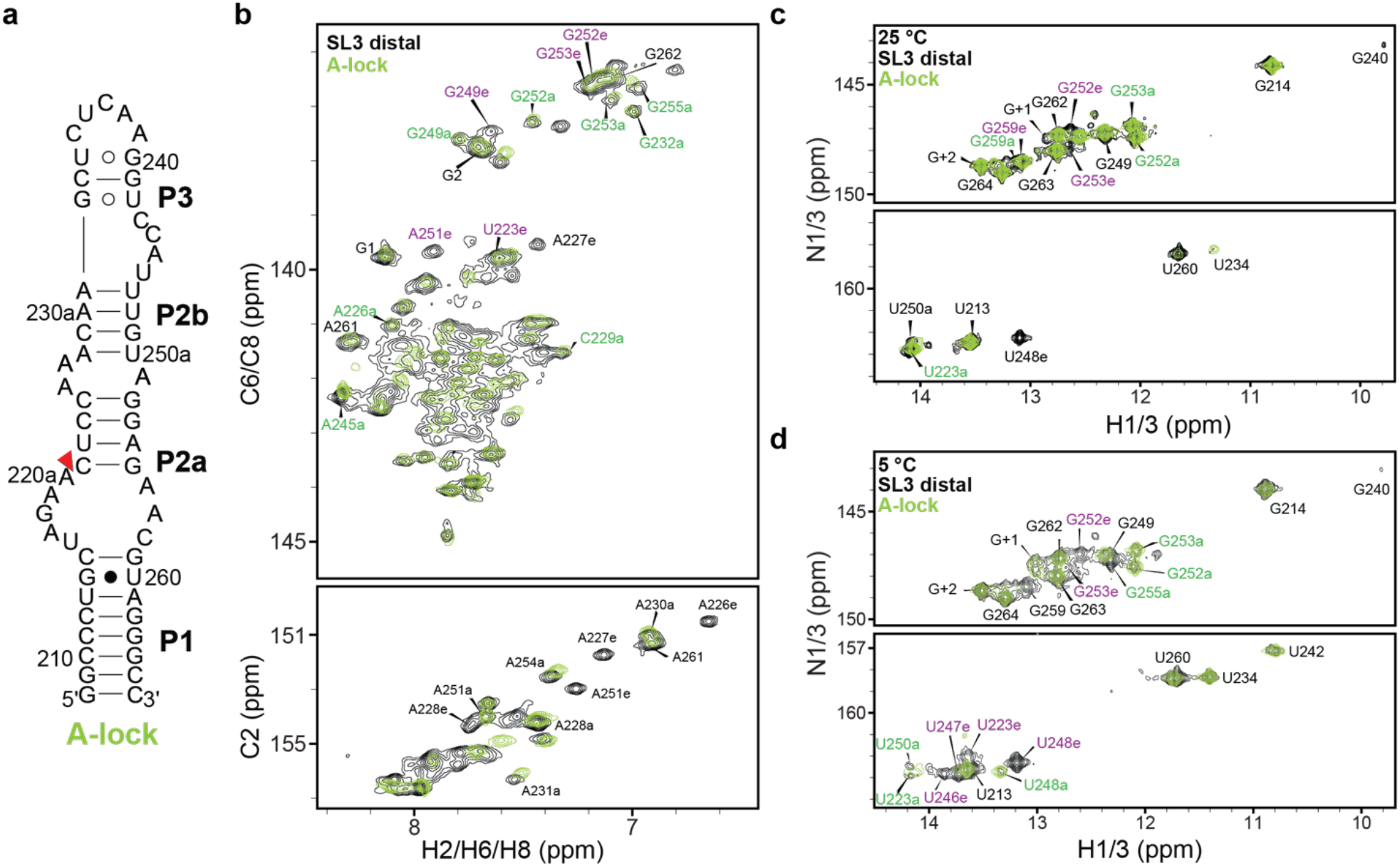
NMR data of SL3 A-lock construct. a) Secondary structure of SL3 A-lock construct with deletion site indicated with a red triangle. b) 2D ^1^H-^13^C HSQC spectra of SL3 A-lock construct (green) overlaid on wild-type SL3 distal construct (black). c-d) Overlay of 2D ^1^H-^15^N HSQC spectra of SL3 A-lock and wild-type SL3 distal constructs. Resonance labels for SL3e state (purple) and SL3a state (green).

**Figure S10.**
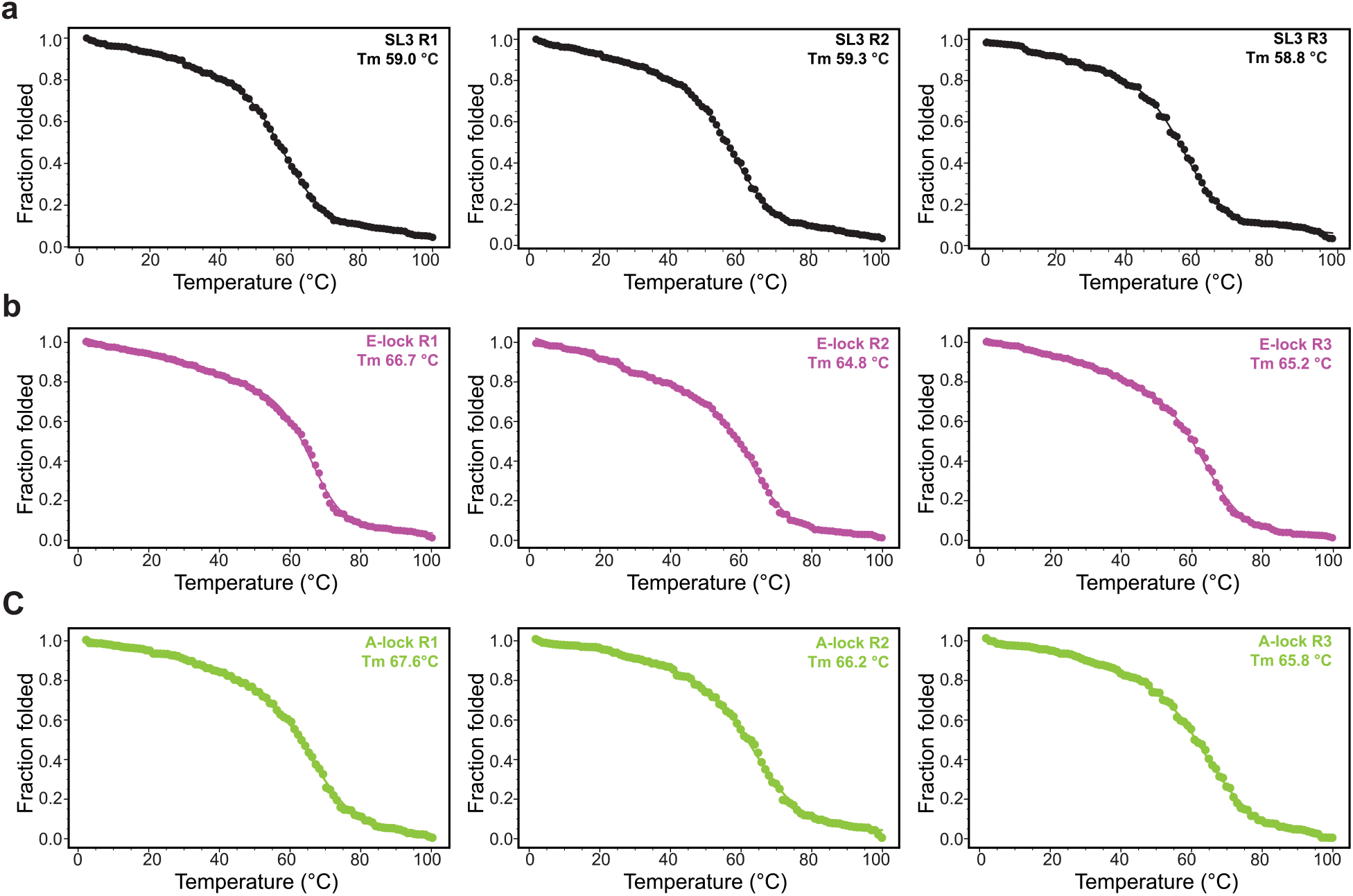
CD thermal melting analysis of SL3 distal, E-lock, and A-lock constructs. R indicates the replicate number for each construct. The data were fitted using Equation 1 (see methods) and plotted with an in-house Python script (https://github.com/ceichhorn2). The experimental data points are shown as circles with the fit line superimposed.

**Figure S11.**
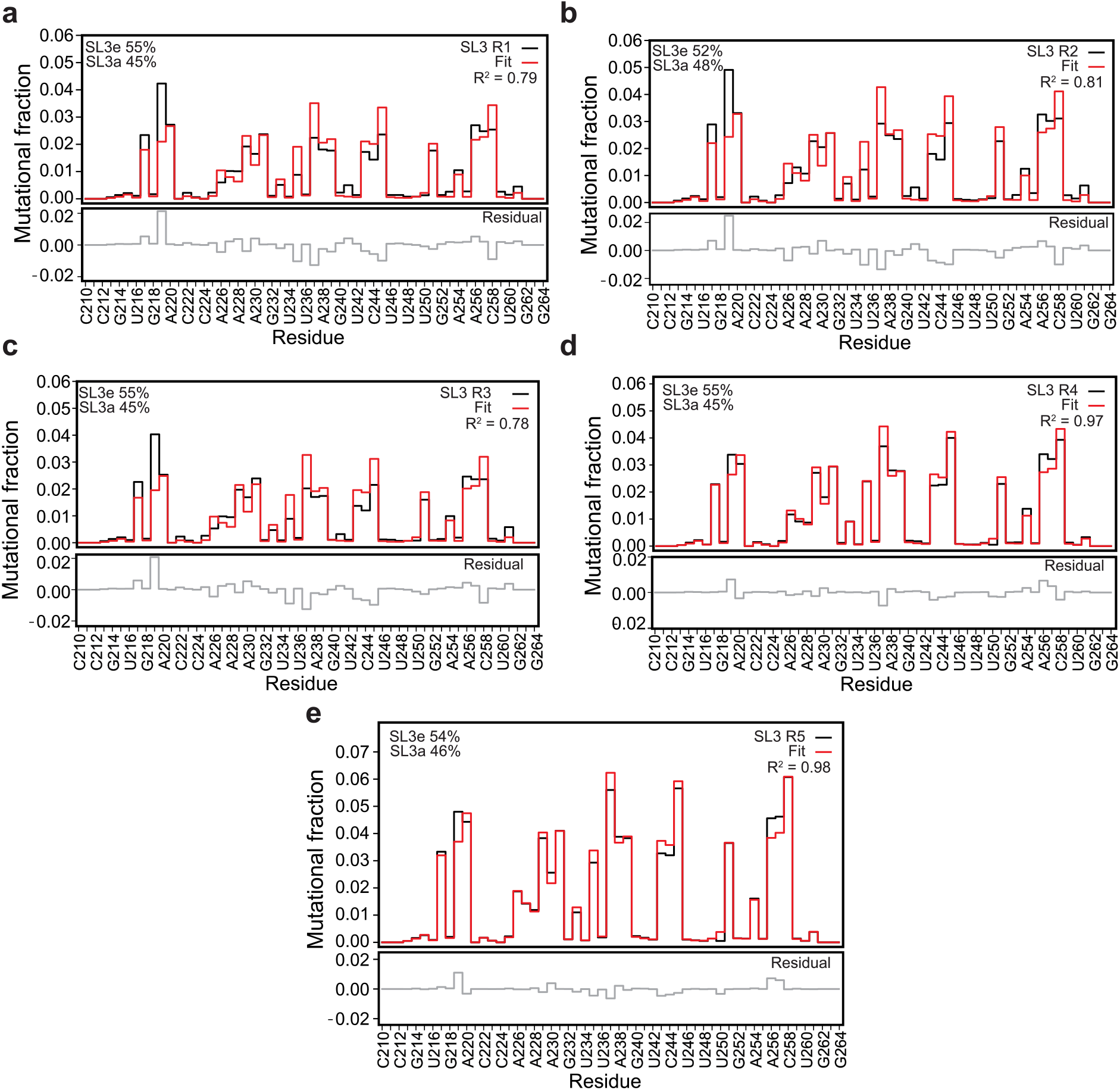
Linear combination analysis of isolated SL3 distal construct. E-lock was used as a fingerprint for the SL3e state, while A-lock was used as a fingerprint for the SL3a state. a-e) Linear combination was performed for each replicate of the SL3 distal construct. The small residual values (grey) and high R^2^ shows a close correlation between SL3 distal (black) and linearly combined E-lock and A-lock (red).

**Figure S12.**
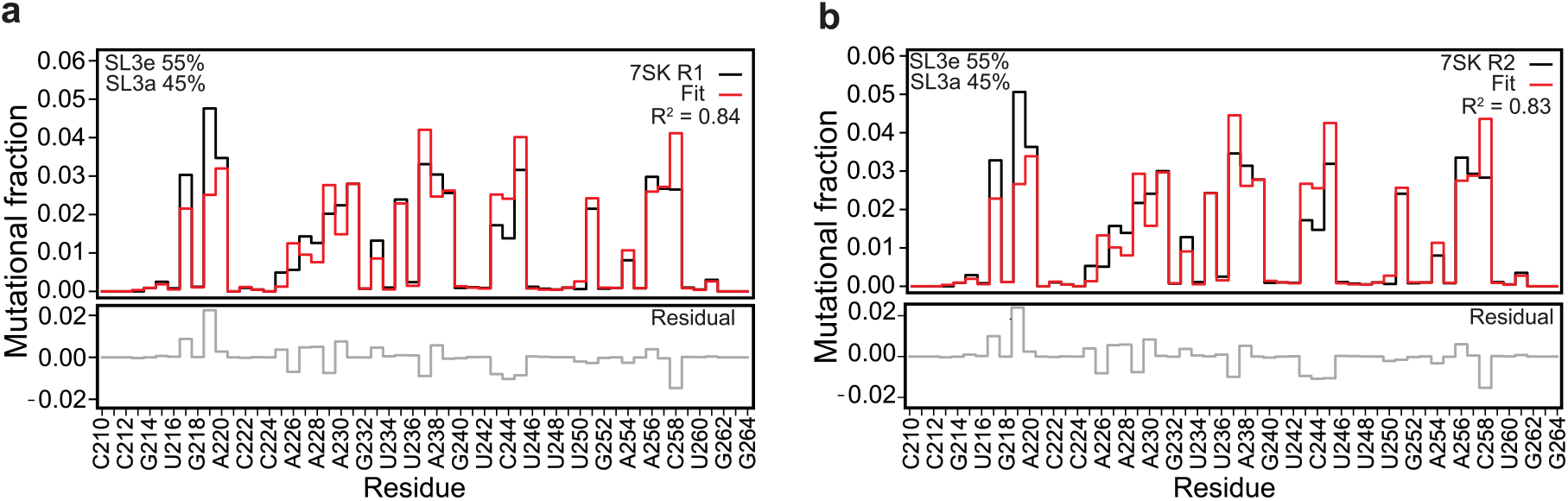
Linear combination analysis of independent replicates of the full-length 7SK RNA (nts 210-264). a) and b) Linear combination analysis was performed from 210-264 nt regions of the 7SK RNA using E-lock and A-lock as fingerprints. The two independent replicates show similar populations and R^2^ correlation.

**Figure S13.**
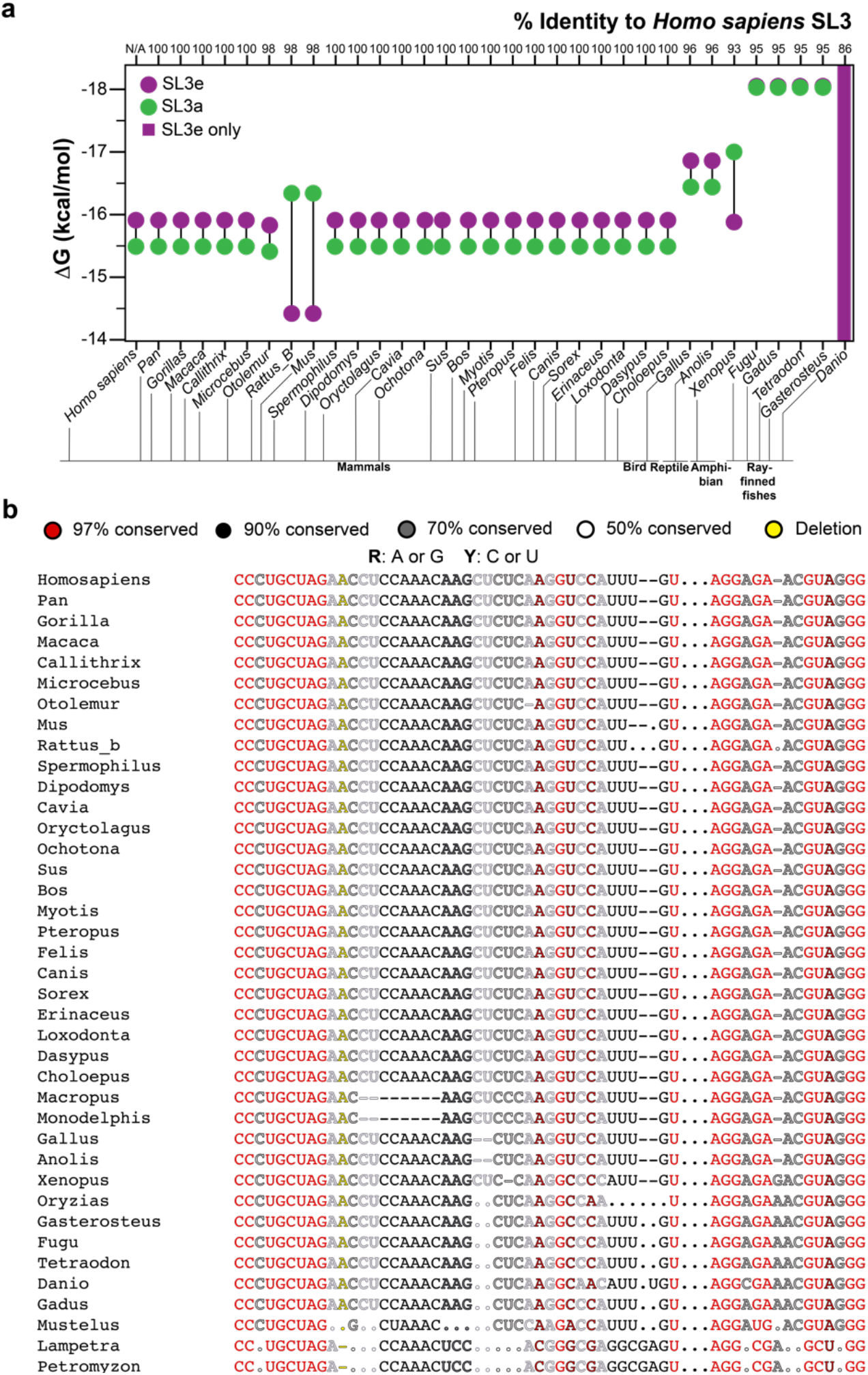
Phylogenetic study of SL3e and SL3a states in vertebrates. a) Predicted ΔG for SL3e and SL3a states from the top five predicted secondary structures of vertebrate sequences. Purple circles or green circles represent SL3e or SL3a states, respectively. Purple bars signify vertebrate sequences where only SL3e state is predicted. We excluded *Oryzias*, *Macropus*, *Monodelphis, Mustelus, Lampetra and Petromyzon* from this analysis because the sequences of these organisms predict neither SL3a nor SL3e states. b) Multiple sequence alignment of vertebrate 7SK RNA SL3 sequences. The sequence has been color-coded based on sequence conservation colors from R2R.

**Figure S14.**
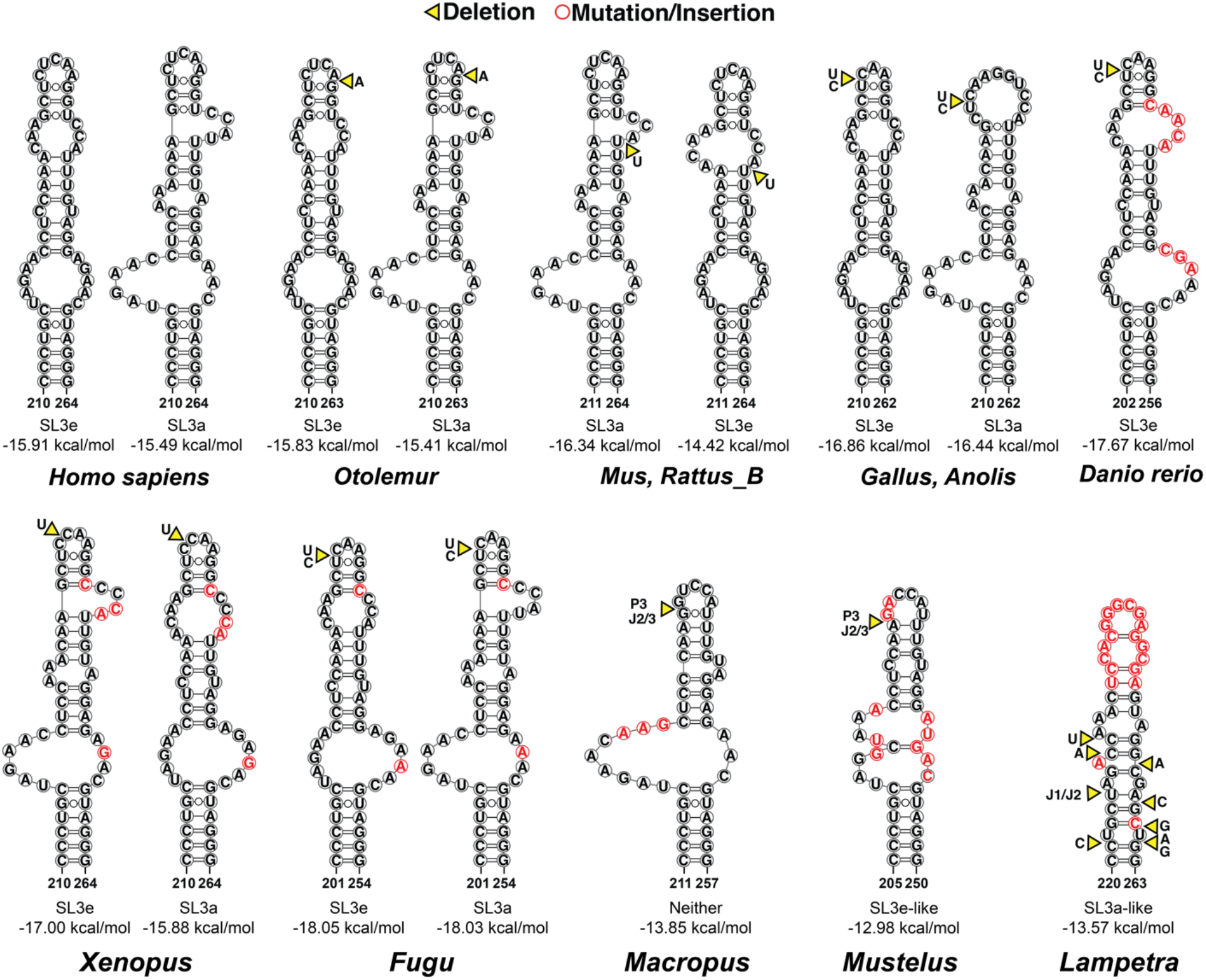
Predicted secondary structures of vertebrate 7SK RNA SL3. Predicted secondary structure of *Homo sapiens* (human) and species with identical sequence, *Otolemur* (greater galago), *Mus musculus, Rattus_B* (mouse/rat), *Gallus gallus* (chicken), *Macropus/Monodelphis* (marsupials), *Xenopus laevis* (African clawed frog), *Fugu* (Takifugu), *Danio rerio* (zebrafish), *Mustelus* (smooth-hound sharks) and *Lampetra* (lampreys). Yellow triangle denotes deleted nucleotides compared to *Homo sapiens* and red circles indicate insertion or mutation. Secondary structure analysis for *Macropu*s, *Mustelus* and *Lampetra* is limited because of less sequence similarity to *Homo sapiens* 7SK RNA SL3.

**Table S1.** Summary of populations from DREEM and linear combination analysis for SL3 distal end constructs and full-length 7SK RNA (nts 210-264).

| Construct | DREEM populations (%) |  | Linear combination population (%) |  |  |  |
| --- | --- | --- | --- | --- | --- | --- |
| **SL3 distal** | **SL3e** | **SL3a** | **SL3e** | **SL3a** | **R<sup>2</sup>** | **Sum of residuals** |
| 1 | 55.0 ± 0.0 | 45.0 ± 0.0 | 55.0 | 45.0 | 0.79 | 0.12 |
| 2 | 49.3 ± 0.5 | 50.7 ± 0.5 | 52.0 | 48.0 | 0.81 | 0.16 |
| 3 | 49.3 ± 0.5 | 50.7 ± 0.5 | 55.0 | 45.0 | 0.78 | 0.14 |
| 4 | 53.0 ± 0.8 | 47.0 ± 0.8 | 55.0 | 45.0 | 0.97 | 0.07 |
| 5 | 54.7 ± 0.5 | 45.3 ± 0.5 | 54.0 | 46.0 | 0.98 | 0.07 |
| Average | 52.3 ± 1.4 | 47.7 ± 1.4 | 54.6 ± 0.8 | 45.4 ± 0.8 | n/a | n/a |
| **7SK RNA** | **SL3e** | **SL3a** | **SL3e** | **SL3a** | **R<sup>2</sup>** | **Sum of residuals** |
| 1 | 53.3 ± 0.5 | 46.7 ± 0.5 | 55.0 | 45.0 | 0.84 | 0.13 |
| 2 | 58.0 ± 0.0 | 42.0 ± 0.0 | 55.0 | 45.0 | 0.83 | 0.15 |
| Average | 55.7 ± 1.6 | 44.3 ± 1.6 | 55.0 ± 0.0 | 45.0 ± 0.0 | n/a | n/a |
| **SL3 E-lock** | **SL3e** | **SL3a** | **SL3e** | **SL3a** | **R<sup>2</sup>** | **Sum of residuals** |
| 1 | 100.0 ± 0.0 | 0.0 ± 0.0 | n/a | n/a | n/a | n/a |
| 2 | 100.0 ± 0.0 | 0.0 ± 0.0 | n/a | n/a | n/a | n/a |
| 3 | 100.0 ± 0.0 | 0.0 ± 0.0 | n/a | n/a | n/a | n/a |
| 4 | 100.0 ± 0.0 | 0.0 ± 0.0 | n/a | n/a | n/a | n/a |
| Average | 100.0 ± 0.0 | 0.0 ± 0.0 | n/a | n/a | n/a | n/a |
| **SL3 A-lock** | **SL3e** | **SL3a** | **SL3e** | **SL3a** | **R<sup>2</sup>** | **Sum of residuals** |
| 1 | 0.0 ± 0.0 | 100.0 ± 0.0 | n/a | n/a | n/a | n/a |
| 2 | 0.0 ± 0.0 | 100.0 ± 0.0 | n/a | n/a | n/a | n/a |
| 3 | 0.0 ± 0.0 | 100.0 ± 0.0 | n/a | n/a | n/a | n/a |
| 4 | 0.0 ± 0.0 | 100.0 ± 0.0 | n/a | n/a | n/a | n/a |
| Average | 0.0 ± 0.0 | 100.0 ± 0.0 | n/a | n/a | n/a | n/a |

DREEM analysis was performed thrice for each replicate. The populations from the three iterations were averaged thrice to give the final populations and standard deviation for each replicate. The final DREEM population for SL3e and SL3a was calculated from the average of all of the replicates. The standard deviation was calculated by taking the sum of the squares of the population from all replicates. Linear combination analysis for each replicate was performed using an in-house Python script (https://github.com/ceichhorn2)

**Table S2.**
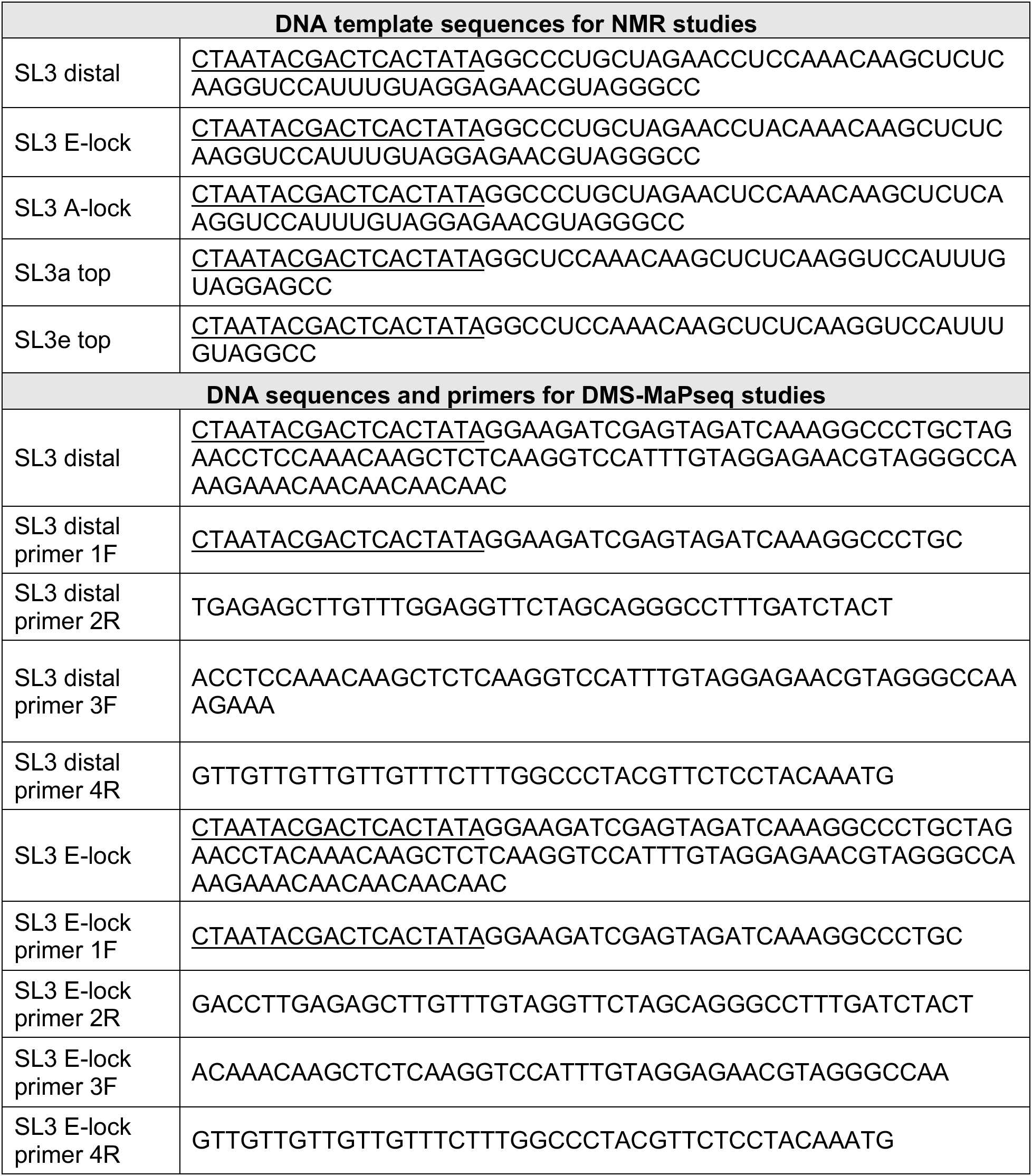
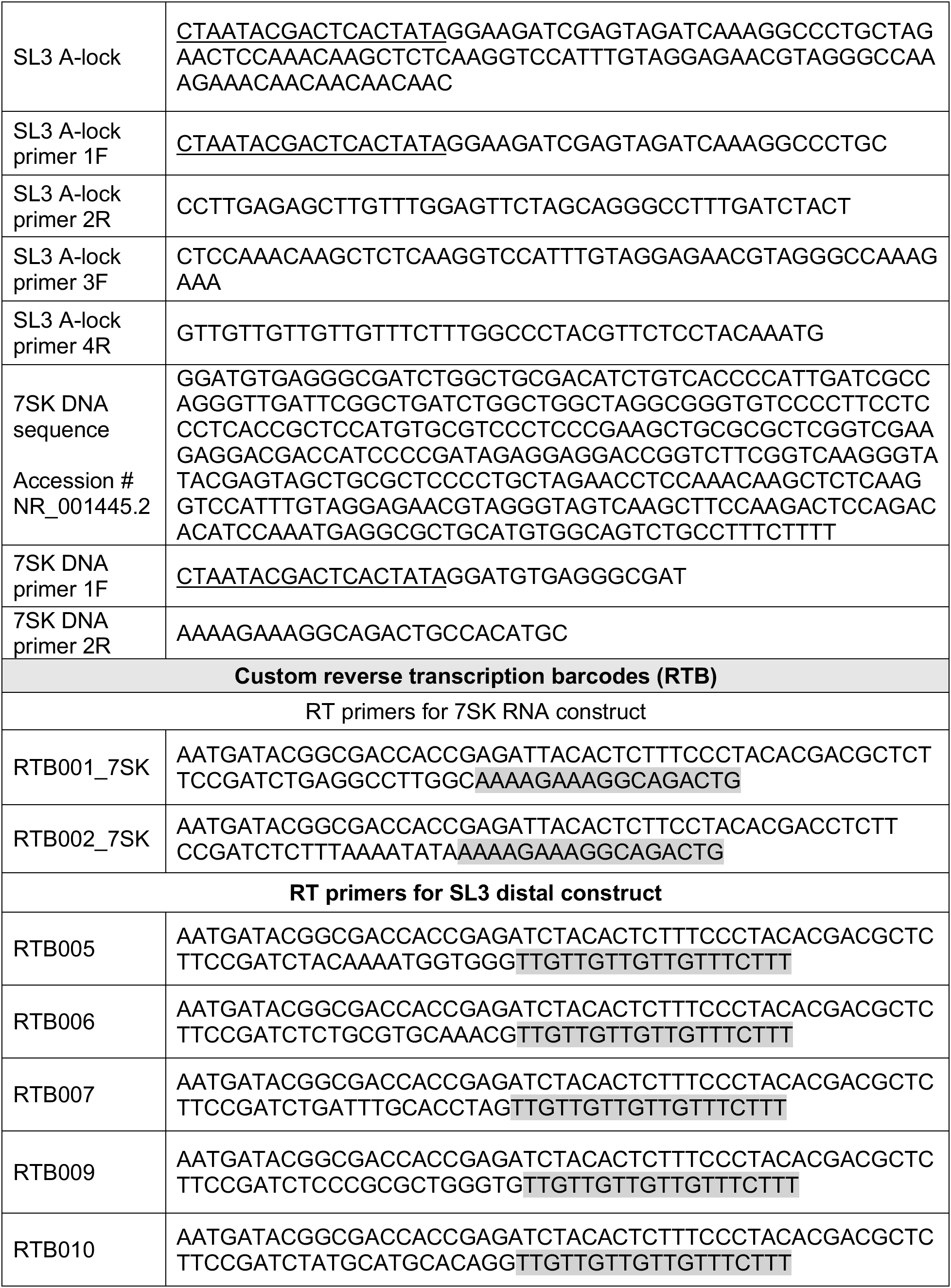
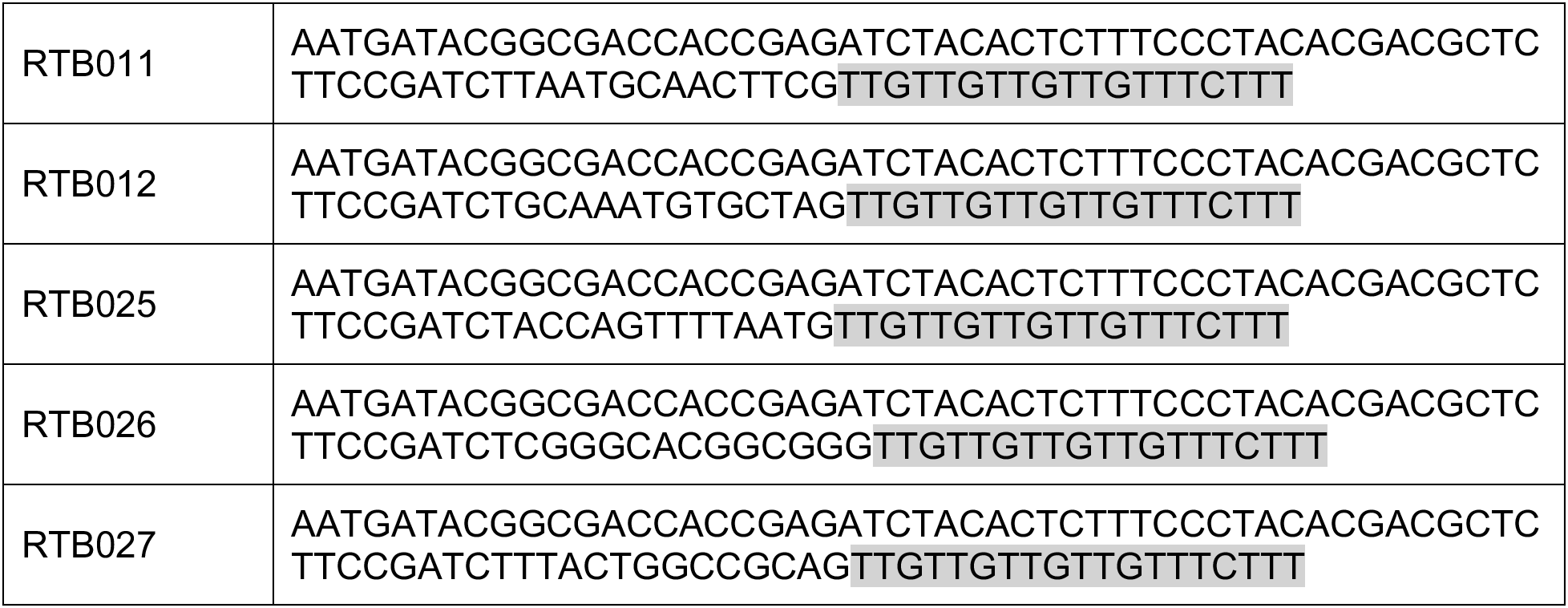
Oligonucleotides and primers used in the study. Underlined nucleotides denote the bacteriophage T7 RNA polymerase promoter sequence for *in vitro* transcription. Highlighted sequences denote reverse transcription barcode sequences complementary to 7SK RNA or SL3 distal constructs for chemical probing experiments.

**Table S3.**
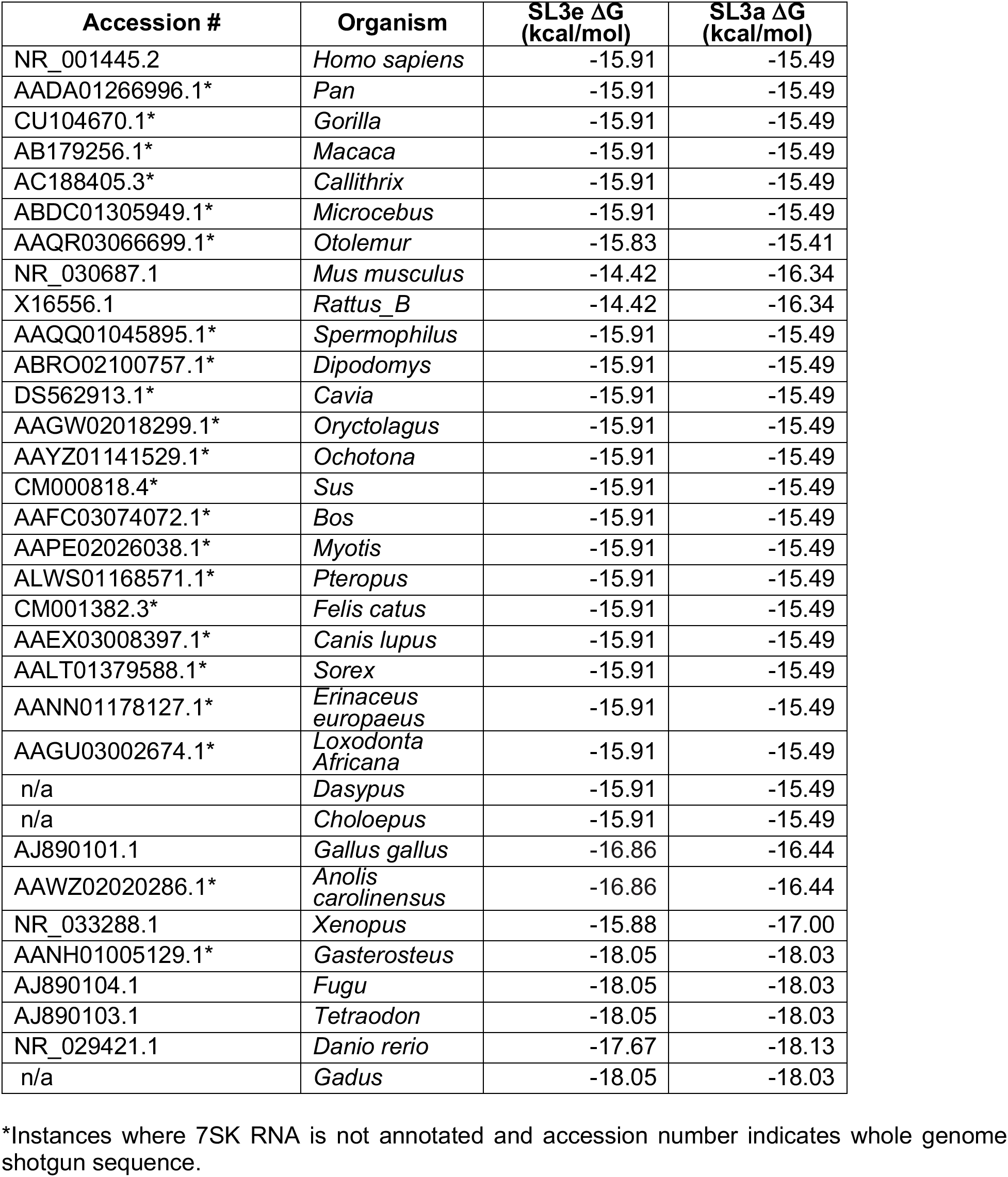
Computed free energy values (ΔG) for vertebrate SL3e and SL3a states using the RNAeval web server.

**Table S4:**
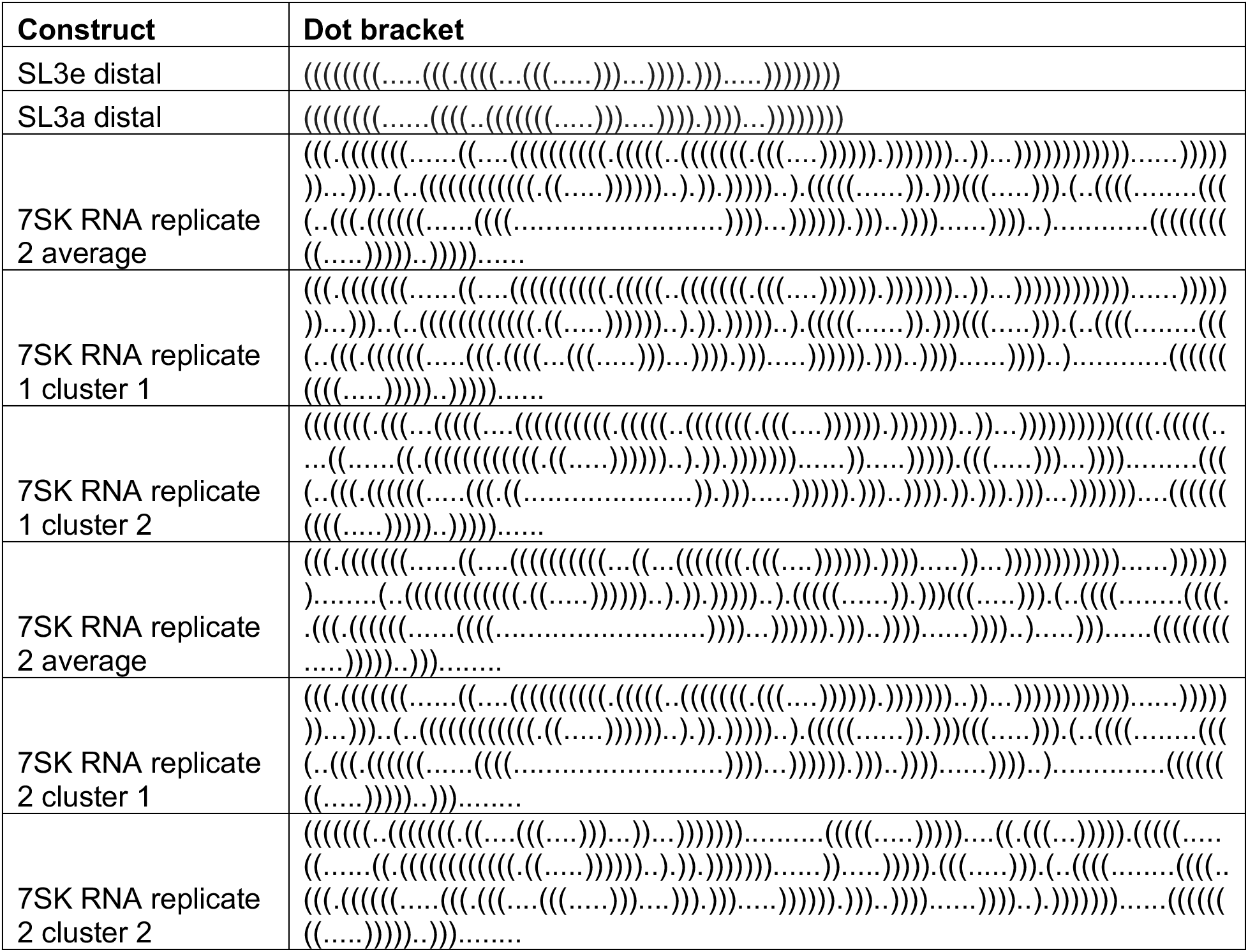
Dot bracket notations of SL3 distal and full-length 7SK RNA secondary structures from DMS-MaPseq studies.

